# Mechanism of *HS5-1* eRNA Regulation of Protocadherin *α* Cluster via R-loop Formation

**DOI:** 10.1101/2021.08.31.458448

**Authors:** Yuxiao Zhou, Siyuan Xu, Qiang Wu

## Abstract

Enhancers generate bidirectional noncoding enhancer RNAs that may regulate gene expression. At present, mechanisms of eRNA functions are not fully understood. Here, we report an antisense eRNA *PEARL* that is transcribed from the protocadherin *α HS5-1* enhancer region. Through loss- and gain-of-function experiments with CRISPR/Cas9 DNA-fragment editing, CRISPRi, and CRISPRa strategies, in conjunction with ChIRP, MeDIP, and DRIP experiments, we find that *PEARL* regulates *Pcdhα* expression by forming local R-loop *in situ* within the *HS5-1* enhancer region to promote long-distance chromatin interactions between distal enhancer and target promoters. These findings have important implications regarding mechanisms by which the *HS5-1* enhancer regulates stochastic *Pcdhα* promoter choice in single cells in the brain.

## Introduction

Enhancers are distal *cis*-acting elements that were originally found to stimulate gene expression in a location-and orientation-independent manner (Banerji et al. 1981). However, recent CRISPR inversion studies showed that enhancers are not orientation-independent *in vivo* at least for those associated with CTCF (CCCTC-binding factor) sites (Guo et al. 2015; Lu et al. 2019). Detailed studies of specific gene loci such as the *β-globin* gene cluster have shed significant insights into our understanding of enhancer function (Collis et al. 1990; Tuan et al. 1992). In addition, transcriptional enhancers can act *in trans* to regulate promoters on homologous chromosomes (Geyer et al. 1990). The enhancer transcription has recently been shown to be a general genome-wide phenomenon (Kim et al. 2010; Ørom et al. 2010) and the transcribed RNAs from active enhancers are known as eRNAs (Kim et al. 2010; Hah et al. 2011; Mousavi et al. 2013). In particular, clusters of strong composite enhancers known as super-enhancers transcribe prevalent eRNAs and activate gene expression programs that often determine cell identities (Loven et al. 2013; Whyte et al. 2013; Xiang et al. 2014; Pefanis et al. 2015). Despite widespread transcription and extensive investigations of eRNAs (Li and Fu 2019; Sartorelli and Lauberth 2020), the exact function of eRNAs remains obscure.

The clustered protocadherin (*cPcdh*) genes are organized into three closely-linked clusters of *α*, *β,* and *γ*. They are stochastically and monoallelically expressed in a cell-specific manner in the brain (Wu and Maniatis 1999; Esumi et al. 2005; Canzio et al. 2019; Jia et al. 2020). The encoded Pcdh proteins are thought to function as neural identity codes to specify tremendous numbers of neuronal connections (Yagi 2012; Canzio et al. 2019; Wu and Jia 2021). The human *Pcdh α* and *γ*, but not *β*, clusters have variable and constant genomic organization, similar to those of the immunoglobulin, T-cell receptor, and UDP-glucuronosyltransferase gene clusters (Wu and Maniatis 1999; Zhang et al. 2004). The variable region of the *Pcdh α* gene cluster contains 13 highly-similar alternate variable exons (*α1-α13*) and two C-type variable exons (*αC1* and *αC2*), each of which is separately spliced to a single set of three downstream constant exons to generate diverse mRNAs (Wu and Maniatis 1999). Each *Pcdhα* alternate variable exon is preceded by a promoter which is flanked by two forward-oriented CTCF sites (Fig. 1A) (Guo et al. 2012; Guo et al. 2015; Canzio et al. 2019; Jia et al. 2020).

**Figure 1.**
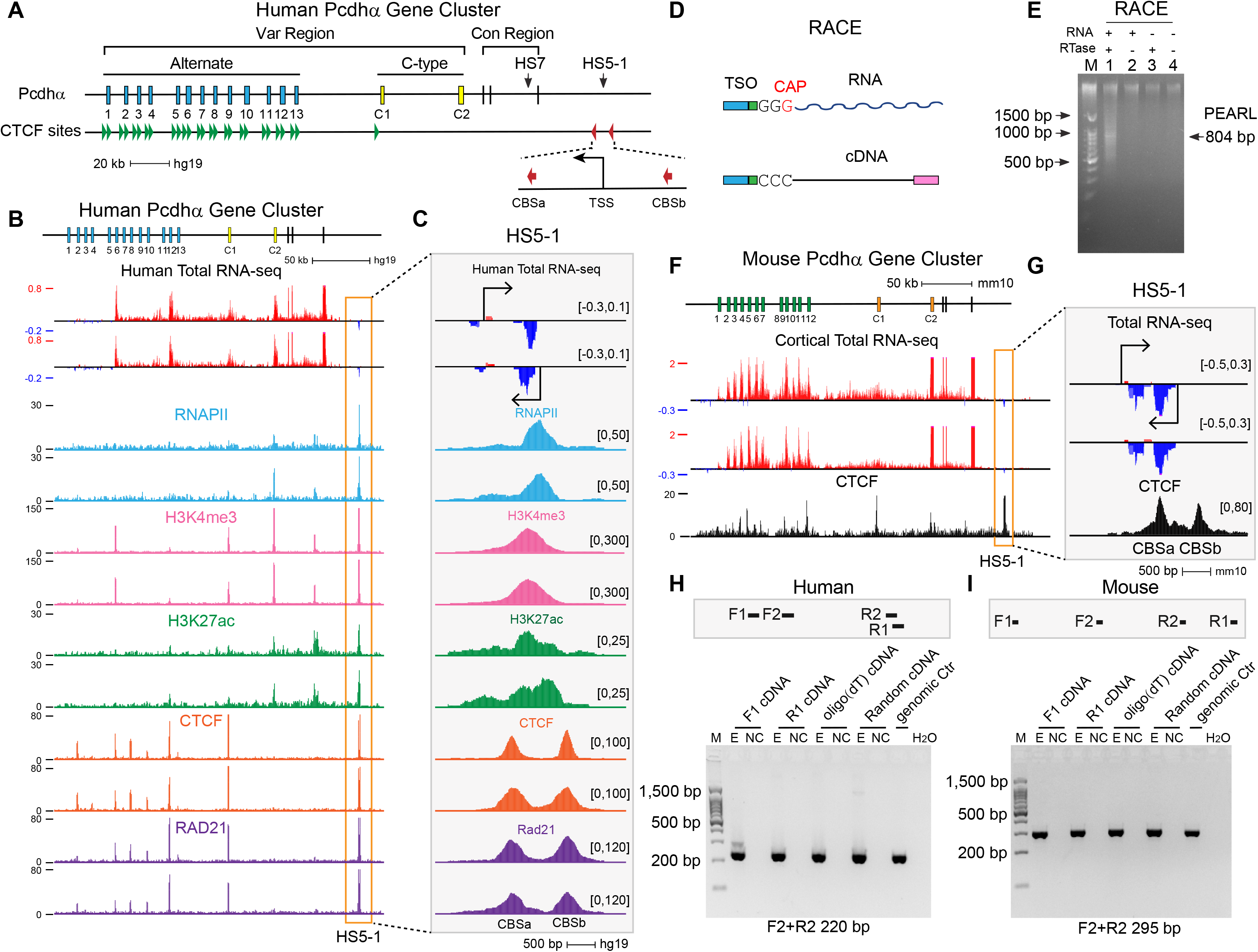
Protocadherin *HS5-1* enhancer produces an antisense eRNA. *(A)* Schematic representation of the human *Pcdhα* cluster. *(B-C)* Shown are total RNA-seq, RNAPII, H3K4me3, H3K27ac, CTCF and Rad21 ChIP-seq in human *Pcdhα* cluster. Total RNA-seq shows transcribed sense (red) and antisense (blue) transcripts. H3K4me3 and H3K27ac mark promoter and enhancer activities, respectively. CTCF and Rad21 each has two binding sites in the *HS5-1* region. *(D)* Schematic of template-switching of the 5’ RACE experiment. *(E)* Agarose gels show *HS5-1* eRNA products. *(F-G)* Mouse cortical total RNA-seq. *(H-I)* Quantitative RT-PCR analyses of the bidirectional eRNAs in HEC-1-B cells and mouse cerebral cortex.

The *Pcdhα* cluster is regulated by a downstream super-enhancer composed of two composite enhancers of *HS5-1* and *HS7* (Ribich et al. 2006). In particular, the *HS5-1* enhancer, flanked by two reverse-oriented CTCF sites, is located at ∼30 kb downstream of the last constant exon (Fig. 1A) (Guo et al. 2012; Guo et al. 2015; Canzio et al. 2019; Jia et al. 2020). CTCF/cohesin-mediated active ‘loop extrusion’ results in long-distance “double clamping” chromatin interactions between the two forward-reverse pairs of convergent CTCF sites and brings the *HS5-1* enhancer in close spatial contact with its target variable promoters to determine the *Pcdhα* promoter choice (Guo et al. 2012; Guo et al. 2015; Canzio et al. 2019; Jia et al. 2020). Specifically, antisense transcription of lncRNA from specific variable antisense promoters, which leads to DNA demethylation by TET enzymes and subsequent recruitment of CTCF proteins, is the key determinant of the *Pcdhα* promoter choice (Canzio et al. 2019). However, the mechanism by which long-distance chromatin interactions between the *HS5-1* enhancer and its target promoters regulate *Pcdhα* promoter choice is not fully understood. Here, we report that an antisense eRNA transcribed from the *HS5-1* enhancer regulates *Pcdhα* gene choice *in cis* through forming R-loop and modifying higher-order chromatin structures.

## Results and Discussion

### Pcdhα HS5-1 antisense eRNA revealed by RACE experiments

Using the model cell line of HEC-1-B (Guo et al. 2015), we first performed strand-specific total RNA-seq experiments, which remove the abundant ribosomal RNAs (Ameur et al. 2011), and found prominent antisense transcripts from the *HS5-1* enhancer region (Fig. 1B,C). These transcripts map to a position enriched of RNAPII (RNA polymerase II), H3K4me3 (histone 3 lysine 4 trimethylation), and H3K27ac (histone 3 lysine 27 acetylation), which are located between the two reverse-oriented CTCF sites (Fig. 1C). To map the exact transcription start site (TSS), we carried out 5′ RACE (rapid amplification of cDNA ends) experiments and found an 804-nt eRNA molecule which we named *PEARL* (*Pcdh* eRNA associated with R-loop formation) (Fig. 1D,E; Supplementary Fig. S1A). Total RNA-seq with mouse cortical tissues also revealed a prominent *HS5-1* antisense transcript (Fig. 1F,G). In addition, sequence analyses showed that these *HS5-1* eRNA transcripts contain no conserved ORF (open reading frame), suggesting that this *HS5-1* eRNA is noncoding. Moreover, in both human HEC-1-B cells and mouse cortical brain tissues, total RNA-seq experiments revealed weaker but detectable *HS5-1* sense transcripts (Fig. 1C,G). To confirm the bidirectional eRNA transcription, we performed quantitative RT-PCR experiments using either a sense or antisense primer as a reverse transcription primer and found that both can produce cDNA (Fig. 1H,I), suggesting that the *HS5-1* enhancer transcription is bidirectional. In addition, we used oligo d(T) as the reverse transcription primer and found that it can produce cDNA, suggesting that the *HS5-1* eRNAs are polyadenylated (Fig. 1H,I). Finally, we quantified eRNA expression levels in model cell lines of HEC-1-B, SK-N-SH, HepG2, and HEK293T and found that *HS5-1* eRNA expression levels correlate with those of *Pcdhα* expression (Supplementary Fig. S1B,C).

### HS5-1 eRNA PEARL TSS deletion affects Pcdhα expression

To investigate the potential function of the *HS5-1* eRNA *PEARL* in regulating *Pcdhα* expression, we specifically deleted the TSS region by CRISPR DNA-fragment editing to avoid the perturbation of the two CBS (CTCF-binding site) elements (Fig. 2A,B), which are known to be essential for *Pcdhα* gene regulation (Guo et al. 2012). We isolated two independent homozygous single-cell CRISPR deletion clones (Supplementary Fig. S1D-F). RNA-seq experiments revealed a significant decrease of *Pcdh α6*, *α12*, *αC1* expression levels upon TSS deletion (Fig. 2C; note that *PcdhαC2* contains no CBS (Guo et al. 2012) and is not regulated by *HS5-1* (Ribich et al. 2006)). Next, we examined histone modifications at the *HS5-1* enhancer and *Pcdhα* alternative promoters to understand the TSS-deletion impact on chromatin states. We first examined enrichment levels of H3K4me3 (Fig. 2D) by ChIP-seq with a specific antibody and found that the TSS deletion results in a significant decrease of H3K4me3 occupancy at the *α6*, *α12*, and *αC1* promoters as well as the *HS5-1* enhancer region (Fig. 2E,H), consistent with the reduction of gene expression of *α6*, *α12* and *αC1* (Fig. 2C). We also performed ChIP-seq experiments with a specific antibody against RNAPII or H3K27ac (Fig. 2I,J) and found that deletion of the eRNA *PEARL* TSS results in a significant reduction in the occupancy of RNAPII and H3K27ac in the *HS5-1* enhancer region (Fig. 2K,L), suggesting that the *HS5-1* enhancer activity is impaired upon eRNA *PEARL* TSS deletion.

**Figure 2.**
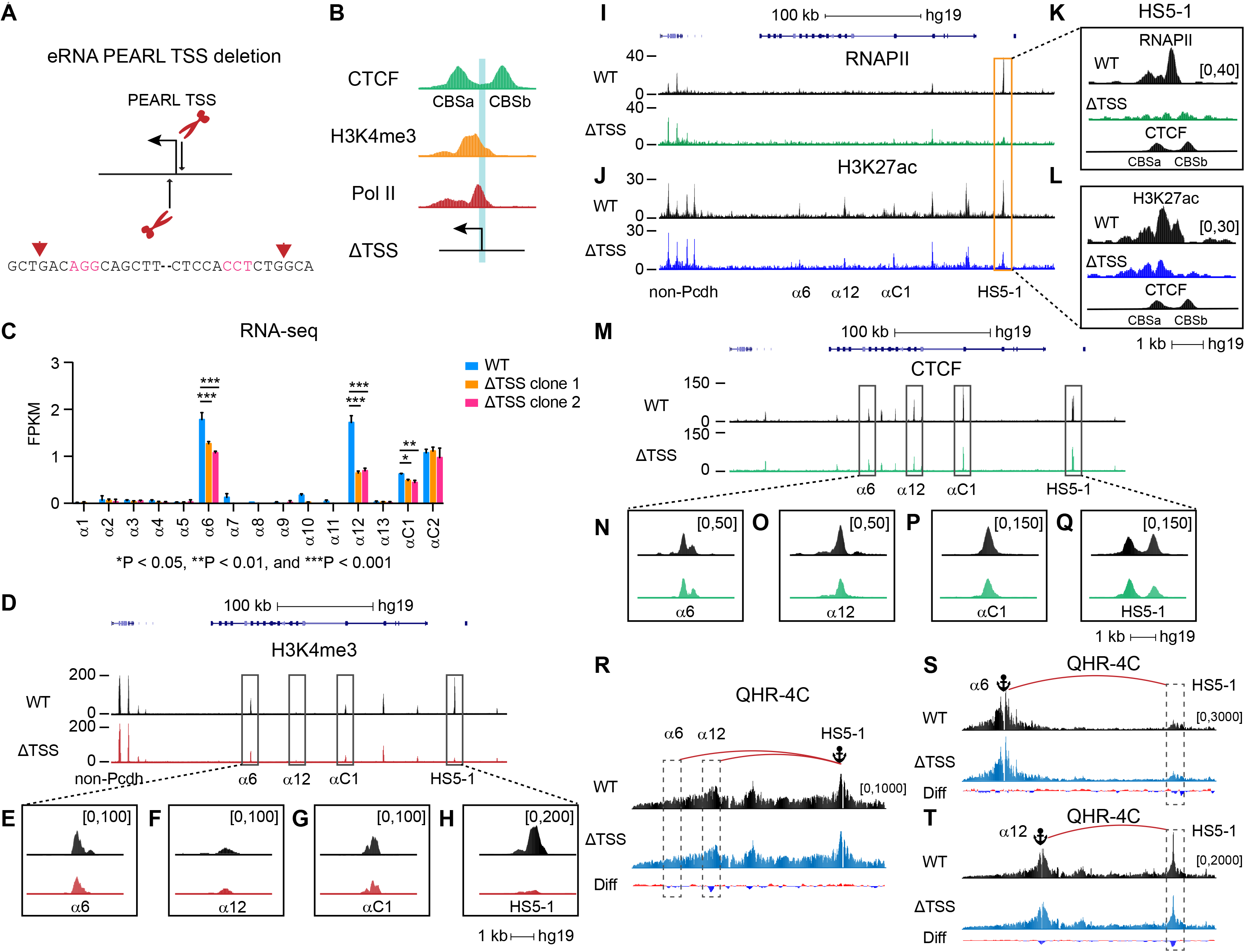
Deletion of *HS5-1* eRNA *PEARL* TSS affects *Pcdhα* expression and chromatin landscape. *(A)* Schematic representation of deleting the *HS5-1* eRNA *PEARL* TSS using the CRISPR/Cas9 system with a pair of sgRNAs. *(B)* Shown is the TSS deletion region relative to H3K4me3 and RNAPII ChIP-seq signals at the *HS5-1* enhancer region. *(C)* RNA-seq for wild-type (WT) and two eRNA TSS deletion single-cell clones in the *Pcdhα* cluster (n = 3). *(D-H)* H3K4me3 ChIP-seq of WT and ΔTSS in the *Pcdhα* cluster. *(I-L)* RNAPII and H3K27ac ChIP-seq of WT and ΔTSS in the *Pcdhα* cluster. *(M-Q)* CTCF ChIP-seq of WT and ΔTSS in the *Pcdhα* cluster. *(R-T)* QHR-4C interaction profiles of WT and ΔTSS with the *HS5-1*, *Pcdhα6,* and *Pcdhα12* as viewpoints (n = 2), represented by anchors.

We next carried out ChIP-seq experiments with a specific antibody against CTCF and found that CTCF occupancy is significantly reduced at the *Pcdh α6*, *α12*, and *αC1* promoters as well as the two sites flanking the *HS5-1* enhancer (Fig. 2M-Q). Quantitative high-resolution chromosome conformation capture copy (QHR-4C) (Jia et al. 2020) with *HS5-1* (Fig. 2R) as a viewpoint revealed a significant decrease of long-distance chromatin interactions between the *HS5-1* enhancer and the *Pcdhα* target genes upon eRNA *PEARL* TSS deletion. Finally, QHR-4C with the alternative *α6* or *α12* promoter as a viewpoint (Fig. 2S,T) confirmed the decreased long-distance chromatin interactions between the distal enhancer and its *Pcdhα* target genes (Fig. 2S,T). Collectively, these data demonstrated that the eRNA *PEARL* TSS is essential for maintaining enhancer activity, mediating chromatin looping, and regulating *Pcdhα* gene expression.

### PEARL regulates Pcdhα gene expression via chromatin looping

We next perturbed the eRNA *PEARL* transcription by inhibiting its initiation, blocking its elongation, or engineering its premature termination by CRISPR epigenetic and genetic methods. CRISPR interference (CRISPRi) can program a KRAB (Krüppel-associated box repressor domain)-fused dCas9 (catalytically dead Cas9) to specific genomic sites to interfere with eRNA transcription without perturbing primary DNA sequences. To this end, dCas9-KRAB is programed by sgRNAs (single-guide RNAs) to a region ranging from 361-bp to 557-bp upstream of the *HS5-1* eRNA *PEARL* TSS to inhibit its transcription initiation (CRISPRi_i) (Fig. 3A) or to a location of 66-bp downstream of TSS to block the RNAPII elongation (CRISPRi_e) (Fig. 3B). There are significant decreases of eRNA expression levels in both cases (Supplementary Fig. S2A). In addition, we inserted a pAS (polyadenylation signal) to cause a premature termination of eRNA transcription (Fig. 3C; Supplementary S2B,C). Two homozygous single-cell CRISPR clones were obtained by screening CRISPR insertion clones and their genotypes were confirmed by PCR and Sanger sequencing (Supplementary Fig. S2D,E).

**Figure 3.**
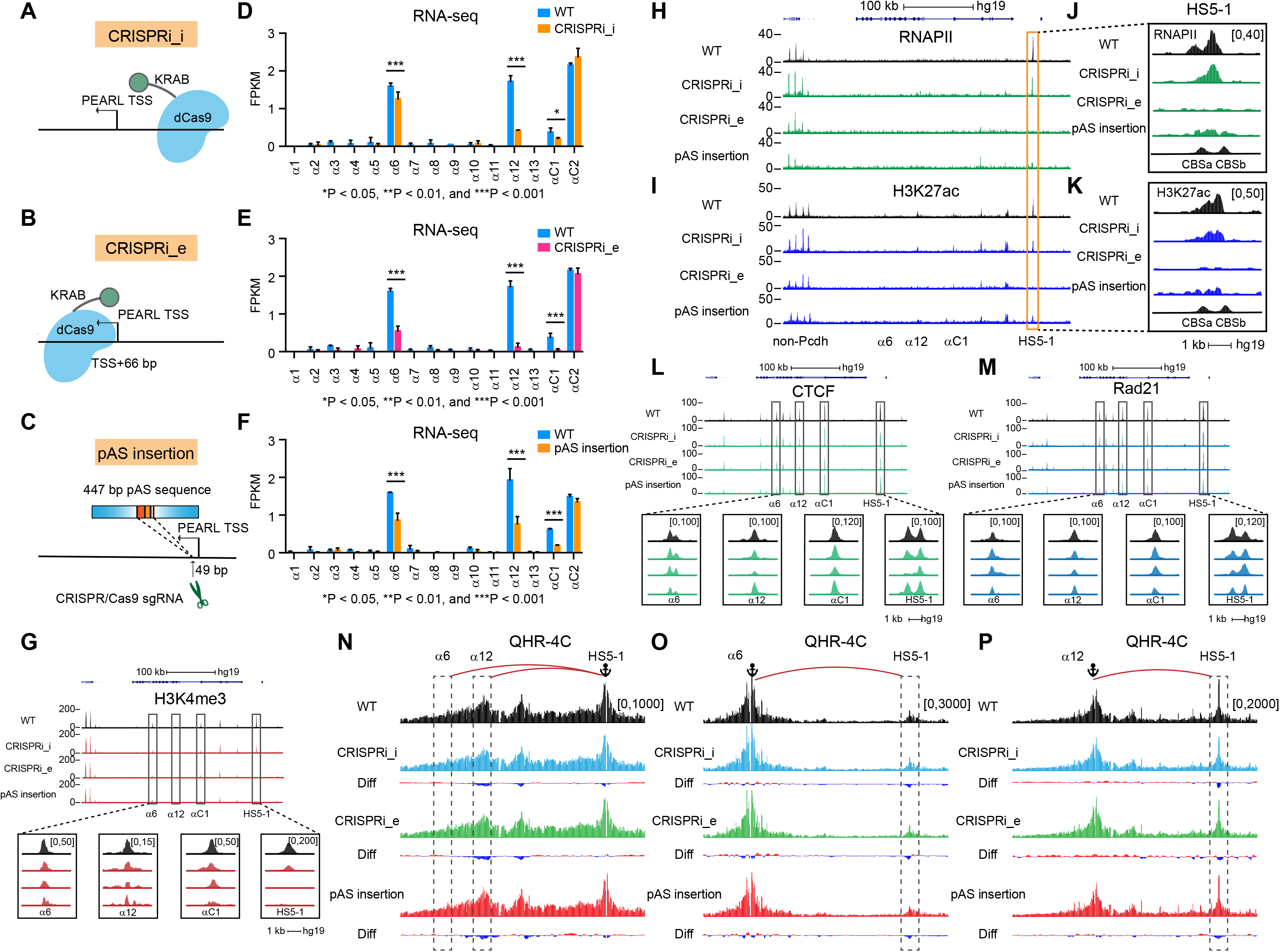
*HS5-1* eRNA *PEARL* affects *Pcdhα* gene expression. *(A-C)* Schematic representation of CRISPR interference (CRISPRi) for perturbation of eRNA *PEARL* transcription initiation (CRISPRi_i) *(A)* or elongation (CRISPRi_e) *(B)*. *P*remature termination of the *HS5-1* eRNA transcription by CRISPR DNA-fragment insertion *(C)*.*(D-F)* Shown are RNA-seq after interfering with the *HS5-1* eRNA *PEARL* transcription initiation, elongation, or termination in the *Pcdhα* cluster (n=3), respectively. *(G)* H3K4me3 ChIP-seq patterns of CRISPRi_i, CRISPRi_e and pAS insertion in the *Pcdhα* cluster. *(H-K)* RNAPII and H3K27ac ChIP-seq of CRISPRi_i, CRISPRi_e, and pAS insertion. *(L-M)* CTCF and Rad21 ChIP-seq of CRISPRi_i, CRISPRi_e, and pAS insertion in the *Pcdhα* cluster. *(N-P)* QHR-4C interaction profiles with the *HS5-1*, *Pcdhα6*, or *Pcdhα12* as a viewpoint (n=2).

RNA-seq experiments revealed a significant decrease of expression levels of *Pcdh α6*, *α12*, and *αC1* in all three methods of perturbing eRNA transcription (Fig. 3D-F). We then examined H3K4me3 levels in the *Pcdhα* cluster after interfering with *PEARL* transcription and found that H3K4me3 enrichments are significantly decreased at the *α6*, *α12*, and *αC1* promoters as well as at the *HS5-1* enhancer (Fig. 3G), consistent with the RNA-seq results (Fig. 3D-F). In addition, RNAPII occupancy and H3K27ac enrichment at the *HS5-1* enhancer are also reduced (Fig. 3H,I). We noted that, compared with perturbing transcription initiation, the RNAPII occupancy and H3K27ac enrichment in the *HS5-1* enhancer region are almost abolished upon blocking eRNA elongation or inserting a premature termination signal (Fig. 3J,K). Next, we designed two sgRNAs to program dCas9 to the locations further downstream of the eRNA *PEARL* TSS (Supplementary Fig. S2F) (One is at 1,158-bp downstream of TSS and the other is at 1,959-bp downstream of TSS). We found that the former reduces the expression levels of both *PEARL* and *Pcdhα* as well as the *HS5-1* enhancer activity, but the latter has no effect, presumably after *HS5-1* eRNA termination (Supplementary Fig. S2G-I). These data suggest that the eRNA *PEARL* is important for *Pcdhα* gene expression.

Previous studies have shown that eRNAs play a role in gene activation by mediating the formation of chromatin looping (Lai et al. 2013; Li et al. 2013; Hsieh et al. 2014; Xiang et al. 2014). In the *Pcdhα* cluster, the stochastic expression of *Pcdhα* depends on the long-distance chromatin interactions between distal enhancer and target variable promoters mediated by the CTCF/cohesin complex. Therefore, we asked whether the eRNA *PEARL* plays a role in CTCF/cohesin-mediated chromatin looping between the distal enhancer and its target variable promoters. To this end, we first measured the enrichments of CTCF and Rad21 by ChIP-seq and found that they were reduced at the *HS5-1* enhancer region (Fig. 3L,M). In addition, they were also reduced at the *Pcdh α6*, *α12*, and *αC1* variable promoters (Fig. 3L,M). We then performed chromosome conformation capture experiments and found that there is a significant decrease of long-distance chromatin interactions between the *HS5-1* enhancer and its target variable promoters (Fig. 3N-P). We concluded that the eRNA *PEARL* is essential for chromatin looping between the *Pcdhα HS5-1* enhancer and its target variable promoters.

### Locally transcribed but not globally overexpressed PEARL regulates Pcdhα expression

To understand the function of *HS5-1* eRNA *PEARL*, we activated eRNA transcription locally through the CRISPR activation (CRISPRa) system with a dCas9-VP160 protein programed to a region upstream of the *HS5-1* eRNA TSS (Supplementary Fig. S3A). We first confirmed the *HS5-1* eRNA CRISPR activation by quantitative RT-PCR (Supplementary Fig. S3B). We then performed RNA-seq and found that *HS5-1* eRNA *PEARL* transcriptional activation results in a significant increase of expression levels of the *Pcdh α6*, *α12*, and *αC1* genes (Supplementary Fig. S3C). Consistently, there is a significant increase of long-distance chromatin interactions between the *HS5-1* enhancer and its target variable promoters (Supplementary Fig. S3D-F).

We next overexpressed the *HS5-1* eRNA *PEARL* globally by a U6 promoter and found, surprisingly, that its overexpression has no effect on *Pcdhα* expression (Supplementary Fig. S3G-I). In addition, its overexpression also has no effect on long-distance chromatin interactions between the *HS5-1* enhancer and its target variable promoters (Supplementary Fig. S3J-L). Finally, we overexpressed the *HS5-1* eRNA *PEARL* in the TSS-deletion CRISPR clones and found that it cannot rescue *Pcdhα* gene expression (Supplementary Fig. S3M-O) nor long-distance chromatin looping between the *HS5-1* enhancer and its target variable promoters (Supplementary Fig. S3P-R). Collectively, we concluded that locally transcribed, but not globally overexpressed, eRNA *PEARL* regulates *Pcdhα* gene expression.

### Local PEARL transcripts form R-loop in situ in the enhancer region

To investigate why locally transcribed, but not globally overexpressed, *PEARL* regulates *Pcdhα* gene expression, we performed chromatin isolation by RNA purification and sequencing (ChIRP-seq) experiments. We synthesized 24 specific 5’-biotin DNA probes directed against the *HS5-1* eRNA *PEARL* transcripts and divided them into odd and even pools to capture with streptavidin magnetic beads (Fig. 4A). We first confirmed that both odd and even probes enrich the *HS5-1* eRNA *PEARL* transcripts (Fig. 4B). We found that there exist powerful specific signals in the *HS5-1* enhancer region with both odd and even probe pools, suggesting that the eRNA *PEARL* is located in the *HS5-1* enhancer region (Fig. 4C,D). As a positive control, the non-coding RNA *NEAT1* is specifically enriched in the *MALAT1* locus *in trans* (Supplementary Fig. S4A) (West et al. 2014). In addition, locked nucleic acid perturbation experiments showed that the *HS5-1* eRNA *PEARL* is required for *Pcdhα* gene expression (Fig. 4E,F). Finally, we performed methylated DNA immunoprecipitation and sequencing experiments (MeDIP-seq) and found that, in contrast to hypermethylation in the *Pcdhα* variable exons (Supplementary Fig. S4B,C), the eRNA promoter within the *HS5-1* enhancer is hypomethylated in both human HEC-1-B cells and mouse cortical brain tissues (Fig. 4G-J). Given that the R-loop structure is often formed by local RNAs in the hypomethylated region (Ginno et al. 2012; Arab et al. 2019; Niehrs and Luke 2020), these MeDIP-seq data suggest that the local eRNA *PEARL* may form R-loop structures in the *HS5-1* region *in situ*.

**Figure 4.**
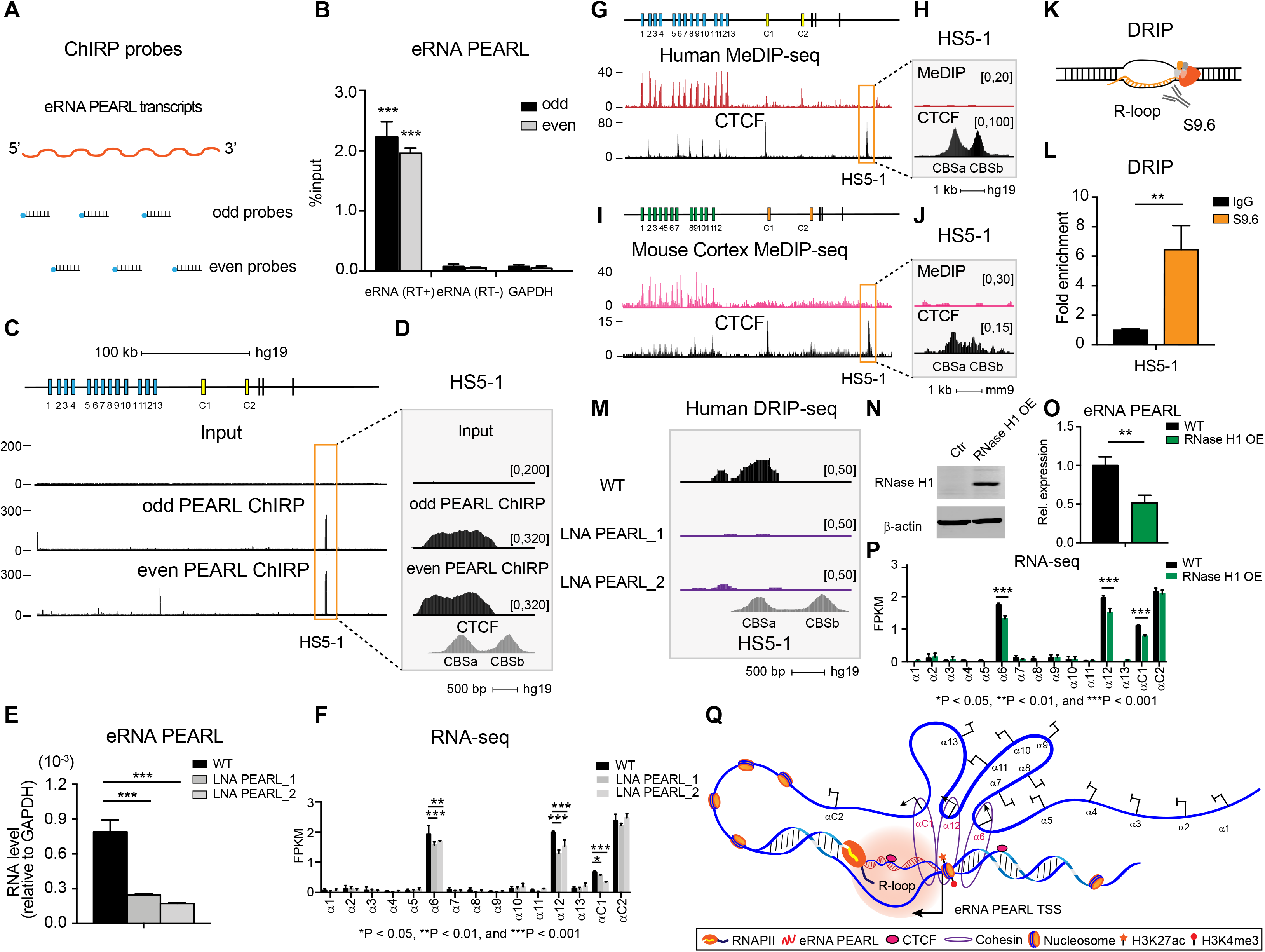
*HS5-1* eRNA *PEARL* forms local R-loop in *HS5-1* enhancer. *(A)* Schematic representation of odd and even probes pools of the ChIRP assay. *(B)* Enrichment of the eRNA *PEARL* after the ChIRP experiments with odd or even probes compared with the housekeeping gene *GAPDH* (n=2). *(C-D)* The *HS5-1* eRNA *PEARL* ChIRP-seq in HEC-1-B cells. *(E)* Shown are the expression profiles of the *HS5-1* eRNA after LNA-mediated silencing of the *HS5-1* eRNA *PEARL* with two functional LNAs, respectively (n=3). *(F)* RNA-seq after LNA perturbation (n=3). *(G-J)* The MeDIP-seq in HEC-1-B cells or mouse cortical tissues. *(K)* Schematic representation of DRIP experiment. *(L)* The R-loop enrichment in the *HS5-1* enhancer region by DRIP-qPCR (n=2). IgG is served as a negative control. *(M)* The R-loop enrichment in the *HS5-1* enhancer by DRIP-seq. *(N)* Western blot of overexpressed RNase H1. OE: overexpression. *(O)* Quantitative analyses of the *HS5-1* eRNA *PEARL* expression after overexpressing RNase H1 (n=3). *(P)* RNA-seq of WT and RNase H1 OE (n=3). *(Q)* The model that *HS5-1* eRNA *PEARL* regulates *Pcdhα* gene expression through R-loop formation.

R-loops or DNA-RNA hybrids are special three-stranded nucleic acid structures that form locally *in vivo* to perform physiological or pathological functions (Crossley et al. 2019; Garcia-Muse and Aguilera 2019; Niehrs and Luke 2020), but whether eRNA in the distal enhancer region regulates gene activation from target promoters via local R-loop *in situ* is not known. To this end, we performed DRIP (DNA-RNA immunoprecipitation) assay with the S9.6 antibody which specifically recognizes DNA-RNA hybrids (Fig. 4K) (Boguslawski et al. 1986; Ginno et al. 2012; Chen et al. 2017; Tan-Wong et al. 2019). We found that the *HS5-1* eRNA is significantly enriched in the S9.6 immunoprecipitates, suggesting that the *HS5-1* eRNA forms R-loop in the enhancer region (Fig. 4L). In addition, locked nucleic acid specific to the eRNA *PEARL* abolishes the *HS5-1* R-loop *in vivo* (Fig. 4M) compared with no effect on the control *NEAT1* R-loop (Supplementary Fig. S4D).

We next overexpressed RNase H1 (Fig. 4N) and found that RNase H1 overexpression results in a significant decrease of the eRNA *PEARL* levels, demonstrating that the eRNA *PEARL* forms DNA-RNA hybrids (Fig. 4O). Moreover, RNase H1 overexpression leads to a significant decrease of expression levels of *Pcdh α6*, *α12*, and *αC1* (Fig. 4P). Furthermore, the long-distance chromatin interactions between the *HS5-1* enhancer and its target variable promoters are also decreased upon RNase H1 overexpression (Supplementary Fig. S4E-G). Finally, we performed ChIRP experiments followed by SDS-PAGE with silver staining and did not find any prominent protein in comparison with the known protein PSF association of *NEAT1* (Supplementary Fig. S4,I) (West et al. 2014; Jiang et al. 2017). Altogether, these data suggest that *HS5-1* eRNA *PEARL* affects *Pcdhα* gene expression through R-loop formation.

### A model for HS5-1 eRNA PEARL function in Pcdhα promoter choice

The ∼60 clustered *Pcdh* genes encode large numbers of cadherin-like cell adhesion proteins that function as neural identify tags in individual cells in the brain (Yagi 2012; Canzio et al. 2019; Wu and Jia 2021). The enormous cell-surface repertoires for neuronal identities are achieved by the combinatorial expression of ∼15 members of the clustered *Pcdh* genes (2 alternate *α* isoforms, 4 *β*, and 4 alternate *γ* isoforms, and 5 C-types *Pcdhs*) (Canzio et al. 2019; Jia et al. 2020). Whereas expression of members of the *Pcdhβγ* clusters is regulated by a super-enhancer downstream of the *Pcdhγ* cluster, members of the *Pcdhα* cluster are regulated by a super-enhancer composed of *HS5-1* and *HS7*. We show here, in mouse and cellular models, that *HS5-1* transcribes a prominent antisense eRNA (Fig. 1; Supplementary Fig. S1) which forms R-loop locally in the *HS5-1* enhancer region (Fig.4; Supplementary Fig.S4) and is required for regulation of distal target promoters through modifying higher-order chromatin architecture. We propose that the eRNA R-loop formation in the *HS5-1* region is essential for establishing the transcription hub (Guo et al. 2012) to activate the chosen promoters of *Pcdh α6*, *α12*, and *αC1* genes (Fig. 4Q). The stochastically chosen *Pcdhα* genes, in conjunction with balanced expression of members of the *Pcdh β* and *γ* clusters, determine individual neuronal identity in the brain.

## Materials and Methods

### Total RNA-seq

Total RNA-seq, which removes the abundant ribosomal RNAs, was performed as previously described (Ameur et al. 2011) with modifications. Briefly, total RNA was extracted from 10^6^ cells of cultured HEC-1-B or postnatal 0-day mouse cortical tissues by TRIzol Reagents (Invitrogen). The ribosomal RNAs were removed from 1 μg of total RNA by incubating with a mixture of biotinylated bait oligonucleotides which are complementary to rRNAs. After reverse transcription of the first-strand cDNA, the second-strand cDNA was synthesized in the directional second strand mix containing dUTP to generate strand-specific cDNA. Libraries were constructed with the NEXTflex Rapid directional RNA-seq Kit (PerkinElmer).

### QHR-4C

Quantitative high-resolution chromosome conformation capture copy (QHR-4C) was performed as recently described (Jia et al. 2020) with modifications. Briefly, as few as 5 × 10^4^ cells were crosslinked in 2% formaldehyde (Thermo) and quenched by 2 M glycine. Cells were lysed twice with 200 μl of cold 4C permeabilization buffer (50 mM Tris-HCl pH 7.5, 150 mM NaCl, 5 mM EDTA, 0.2% SDS, 0.5% NP-40, 1% Triton X-100, and 1 × protease inhibitors) for 10 min at 4°C. Cells were incubated in 73 μl of nuclease-free water, 10 μl of 10 × *Dpn*II buffer, and 2.5 μl of 10% SDS at 37°C with constant shaking at 900 rpm. After 1 h, 12.5 μl of 20% Triton X-100 were then added to the resuspended solution, and the solution continued incubating for 1 h at 37°C with constant shaking at 900 rpm. Two microliters of *Dpn*II (NEB) were added into the reaction. The digested nuclei were ligated with 1 μl of T4 DNA ligase (NEB) in 100 μl of 1 × T4 ligation buffer for 24 h at 16°C. The DNA was sonicated using a Bioruptor system to obtain DNA fragments ranging from 200∼600 bp. A linear amplification step with a specific 5’ biotin-tagged primer complementary to the viewpoint was performed in 100 μl of PCR system. The biotinylated ssDNA was ligated with 0.1 μM adapters in 15 μl of ligation buffer for 24 h at 16°C. After ligation, 8 μl of Streptavidin Magnetic Beads (Invitrogen) were added to capture biotinylated ssDNA. QHR-4C libraries were purified by the High-Pure PCR Product Purification kit (Roche).

### ChIRP-seq

Chromatin Isolation by RNA Purification (ChIRP) was performed as previously described (Chu et al. 2011) with modifications. Briefly, 2 × 10^7^ cells were used for each ChIRP experiment. HEC-1-B cells were cross-linked with 1% glutaraldehyde (Sigma-Aldrich) and quenched with 1 ml of 1.25 M glycine for 5 min. The pellet was resuspended with the specified volume of ChIRP lysis buffer (50 mM Tris-HCl pH 7.0, 10 mM EDTA, 1% SDS, 1 × protease inhibitors) according to its weight. The cell lysate was sonicated using the Bioruptor system to obtain DNA fragments ranging from 100∼500 bp. After centrifuging, 2 ml of ChIRP hybridization buffer (50 mM Tris-HCl pH 7.0, 750 mM NaCl, 1 mM EDTA, 1% SDS, 15% formamide, 1 × protease inhibitors) were added into the cell lysate. Pools of biotinylated odd or even *HS5-1* eRNA *PEARL* probes were added. The captured eRNA chromatin complex was enriched with 100 μl of precleared Streptavidin beads. The beads were purified for DNA library construction with the NEXTflex Rapid DNA-seq Kit (PerkinElmer).

### DRIP-seq

DNA-RNA immunoprecipitation (DRIP) was performed as previously described (Ginno et al. 2012) with modifications. Briefly, 10^7^ HEC-1-B cells were collected and incubated overnight at 55°C in 500 μl of lysis buffer (10 mM Tris-HCl pH 8.0, 1 mM EDTA, 0.5% SDS) with 5.5 μl of 10 mg/ml proteinase K overnight at 37°C. The DNA was purified by phenol-chloroform extraction and ethanol precipitation. The precipitated DNA was dissolved in 200 μl of 10 mM Tris-HCl (pH 7.5) and fragmented by *Xba*I, *Hin*dIII, *Ssp*I, and *Bsr*GI (NEB) overnight at 37°C. Ten micrograms of fragmented DNA were incubated in 500 μl of immunoprecipitation buffer (15 mM Tris-HCl pH 7.5, 1 mM EDTA) with 10 μl of S9.6 antibody (Millipore) rotating overnight at 4°C to capture R-loop *in vivo*. The captured DNA-RNA hybrids were enriched by protein A-agarose beads (Millipore). The purified DNA was used to construct a library with the NEXTflex Rapid DNA-seq Kit (PerkinElmer).

## Acknowledgments

We thank all members of the Wu lab for helpful discussion. This work was supported by grants to Q.W. from the National Natural Science Foundation of China (91940303 and 31630039), the Ministry of Science and Technology of China (2017YFA0504203 and 2018YFC1004504), and the Science and Technology Commission of Shanghai Municipality (19JC1412500).

## Author Contributions

Q.W. conceived and supervised the project. Y.Z. performed most experiments and analyzed data. S.X. helped experiments. Y.Z. and Q.W. wrote the manuscript.

## List of Supplemental Material

### Supplemental Materials and Methods

#### Cell culture

HEC-1-B cells were cultured in modified Eagle’s medium (HyClone) with 10% FBS (ExCell Bio), 1% penicillin-streptomycin solution (Gibco), 2 mM GlutaMAX (Gibco), and 1 mM sodium pyruvate (Gibco). SK-N-SH cells were cultured in MEM (HyClone) supplemented with 10% FBS (Gibco), 2 mM GlutaMAX (Gibco), 1 mM sodium pyruvate (Gibco), and 1% penicillin-streptomycin (Gibco). HepG2 and HEK293T cells were cultured in Dulbecco’s modified Eagle’s medium (HyClone) supplemented with 10% FBS (ExCell Bio) and 1% penicillin-streptomycin (Gibco). All cells were usually cultured at 37°C in the incubator supplied with 5% (v/v) of CO_2_.

#### Plasmid construction

The plasmids of sgRNAs for CRISPR DNA-fragment editing and CRISPR homologous recombination were constructed as previously described (Li et al. 2015; Shou et al. 2018). Briefly, pairs of complementary oligonucleotides were annealed to generate sgRNA constructs with 5′ overhangs of ‘ACCG’ and ‘AAAC’. The annealed dsDNA was cloned into a *Bsa*I-linearized pGL3 vector with U6 promoter. To insert 447-bp pAS (polyadenylation signal) sequences into locations downstream of the *HS5-1* eRNA transcription start site, the plasmid with the inserted sequences and about 1,000 bp flanking homologous arms was used for CRISPR homologous recombination. Two homologous arms, the 447-bp pAS sequences, and the linearized pUC19 vector were jointed together with 20-bp overlapping sequences using a One-Step Cloning recombination system (Vazyme).

The plasmids of sgRNAs for CRISPRi and CRISPRa were constructed to inhibit or activate *HS5-1* eRNA transcription. Pairs of complementary oligonucleotides with 5′ overhangs of ‘CACC’ and ‘AAAC’ were annealed to generate sgRNA constructs. The annealed dsDNA was cloned into a linearized lentivirus sgRNA vector (Addgene) digested with *Bsm*BI. The dCas9-VP160 plasmid was generated by substituting the VP64 sequence of dCas9-VP64 lentivirus backbone plasmid (Vazyme). The VP160 sequence was amplified and cloned into the linearized dCas9-VP64 vector digested by *Bam*HI and *Bsr*GI.

The plasmid for overexpressing eRNA *PEARL* was cloned into a linearized pGL3 vector. The coding sequence of RNase H1 was amplified by PCR from human cDNA and cloned into a linearized pLVX vector digested by *Eco*RI and *Bam*HI. Primers for plasmid construction are listed in Supplementary Table S1. All recombinant plasmids were confirmed by Sanger sequencing.

#### Genetic deletion of *HS5-1* eRNA *PEARL* TSS using CRISPR DNA-fragment editing

CRISPR/Cas9 DNA-fragment deletion was performed as recently described (Li et al. 2015; Shou et al. 2018). Cas9 protein and a pair of sgRNAs were used to delete approximately 300-bp region around *HS5-1* eRNA *PEARL* TSS. The CRISPR DNA-fragment editing components, consisting of Cas9-expressing plasmid and dual sgRNA-expressing plasmids, were co-transfected in a 6-well plate when cells reached ∼60% confluency using Lipofectamine 3000 reagents (Invitrogen) according to the protocol recommended by the manufacturer. After 48 h, puromycin was added to the culture medium at the final concentration of 2 μg/ml. After four days of puromycin selection, cultured cell mixture was screened for the desired deletion band. Then, the cells were diluted and seeded into 96-well culture plates. The single-cell CRISPR clones were cultured for about 2 weeks and the colonies were marked under a microscopy. These colonies continued to be cultured in fresh media to ∼90% confluency. The single-cell CRISPR clones were collected and amplified using primers spanning the deleted sequences. The deletion homozygous single-cell CRISPR clones were confirmed by Sanger sequencing. Two homozygous TSS deletion clones were obtained from a total of 179 single-cell CRISPR clones. Primers for CRISPR DNA-fragment editing are listed in Supplementary Table S1.

#### Lentivirus packaging for establishing stable cell lines

High-titer lentivirus transduction in HEC-1-B cells is an efficient way to establish stable cell lines for inhibiting *HS5-1* eRNA transcription (CRISPRi) or activating *HS5-1* eRNA transcription (CRISPRa). The mixture of the lentivirus transfer plasmid, packaging helper plasmid (psPAX2, Addgene), and an envelope helper plasmid (pMD2.G, Addgene) was co-transfected into HEK293T cells using Lipofectamine 3000 reagents (Invitrogen) to produce lentivirus particles. After 24 h, the medium was replaced with fresh complete medium supplemented with 30% FBS. After transient transfection for 48 h and 72 h, lentivirus particles were harvested from HEK293T cell supernatant and prepared for lentivirus transduction in HEC-1-B cells to establish stable cell lines.

#### Interfering with *HS5-1* eRNA *PEARL* transcription initiation (CRISPRi_i)

CRISPR interference of *HS5-1* eRNA transcription initiation was performed as previously described (Gilbert et al. 2013; Larson et al. 2013) with modifications. The dCas9-KRAB fusions, programed by sgRNAs to the promoter region of *HS5-1* eRNA, repress its transcription initiation, leading to the H3 lysine 9 trimethylation (H3K9me3) deposition in the transcription initiation region (Thakore et al. 2015). The CRISPR-based loss-of-function assay was used to perturb the *HS5-1* eRNA transcription initiation in HEC-1-B cells. These sgRNAs were designed to target to a location upstream of the *HS5-1* eRNA TSS to interfere with eRNA transcription initiation. Non-targeting sgRNA vector was used as a negative control. Briefly, HEC-1-B cells were transducted by dCas9-KRAB lentivirus particles for 24 h when cells reached ∼60% confluency and selected by 20 μg/ml blasticidin for 10 days to obtain stable dCas9-KRAB-expressing cell line. The stable dCas9-KRAB-expressing cells were transducted by lentivirus particles for 24 h to stably express sgRNAs and selected by 2 μg/ml puromycin for 4 days. After puromycin selection, both dCas9-KRAB and sgRNAs were stably expressed in HEC-1-B cells. Primers for CRISPRi_i are listed in Supplementary Table S1.

#### Interfering with *HS5-1* eRNA *PEARL* transcription elongation (CRISPRi_e)

CRISPR interference of *HS5-1* eRNA transcription elongation was performed as previously described (Larson et al. 2013) with modifications. *HS5-1* eRNA transcription elongation was perturbed by programing dCas9-KRAB into locations downstream of *HS5-1* eRNA TSS. The sgRNAs were designed to target to a location downstream of the *HS5-1* eRNA TSS to perturb *HS5-1* eRNA transcription elongation. Non-targeting sgRNA vector was used as a negative control. Briefly, HEC-1-B cells were transducted by dCas9-KRAB lentivirus particles when cells reached ∼60% confluency to obtain stable dCas9-KRAB-expressing cells. Then, the stable dCas9-KRAB-expressing cells were transducted by sgRNA lentivirus particles for 24 h and selected by 2 μg/ml puromycin for 4 days. After puromycin selection, both dCas9-KRAB and sgRNAs were stably expressed in HEC-1-B cells. Primers for CRISPRi_e are listed in Supplementary Table S1.

#### Insertion of premature termination sites using CRISPR homologous recombination

CRISPR/Cas9-mediated homologous recombination was performed as recently described (Li et al. 2015; Shou et al. 2018). A DNA fragment of 447-bp with polyadenylation signal (pAS) sequences was inserted at specific sites using CRISPR homologous recombination to cause premature transcription termination of the *HS5-1* eRNA *PEARL*. The donor plasmid includes 447-bp polyadenylation signal sequences with two ∼1,000 bp flanking homologous arms. This 447-bp transcription termination signal is comprised of a 273-bp PGK pAS, a 125-bp SV40 pAS and a 49-bp synthetic pAS sequences (Joung et al. 2017). Cas9-expressing plasmid, a sgRNA-expressing plasmid and a homologous recombination donor were co-transfected in a 6-well plate when cells reached ∼60% confluency using Lipofectamine 3000 reagents (Invitrogen). Puromycin was added to the culture medium at the final concentration of 2 μg/ml after 48 h. Four days later, the cultured cell mixture was screened for desired insertion bands. Then, the cells were diluted and seeded into 96-well culture plates. The single-cell CRISPR clones were cultured for about 2 additional weeks and the colonies were marked under a microscopy. These colonies were cultured in fresh media for several days to ∼90% confluency. The single-cell CRISPR clones were collected and subsequently amplified using primers spanning homologous arms. The insertion homozygous single-cell CRISPR clones were confirmed by Sanger sequencing. Two homozygous insertion clones were obtained from a total of 698 single-cell CRISPR clones. Primers for CRISPR homologous recombination are listed in Supplementary Table S1.

#### Activating *HS5-1* eRNA *PEARL* Transcription (CRISPRa)

CRISPR activation of *HS5-1* eRNA *PEARL* transcription was performed as previously described (Maeder et al. 2013; Perez-Pinera et al. 2013) with modifications. The CRISPR-based gain-of-function assay was used to activate *HS5-1* eRNA transcription in HEC-1-B cells. These sgRNAs were designed to target to a location upstream of the *HS5-1* eRNA *PEARL* TSS to activate eRNA transcription. Non-targeting sgRNA was used as a negative control. Briefly, HEC-1-B cells were transducted by dCas9-VP160 lentivirus particles for 24 h when cells reached ∼60% confluency and selected by 20 μg/ml blasticidin for 10 days to obtain stable dCas9-VP160-expressing cells. The stable dCas9-VP160-expressing cells were transducted by lentivirus particles for 24 h to express sgRNAs and selected by 2 μg/ml puromycin for 4 days. After puromycin selection, both dCas9-VP160 and sgRNAs were stably expressed in HEC-1-B cells. Primers for CRISPRa are listed in Supplementary Table S1.

#### LNA-ASO knockdown

Cells were transfected with locked nucleic acid antisense oligonucleotides (LNA ASOs) as described previously (Wahlestedt et al. 2000) with modifications. Briefly, the LNA-ASOs complementary to the sequence of *HS5-1* eRNA *PEARL* were designed to knockdown *HS5-1* eRNA *PEARL* transcripts, and a non-targeting LNA-ASO scramble was used as a negative control. 5 × 10^5^ cells per well were seeded in a 6-well plate on the previous day of transfection. LNA-ASOs were resuspended in nuclease-free water and transfected to HEC-1-B cells when cells reached ∼60% confluency with Lipofectamine 3000 transfection reagents (Invitrogen). The sequences of LNA ASOs are listed in Supplementary Table S1. All LNA-ASO experiments were performed with at least three biological replicates.

#### RACE

Rapid amplification of cDNA ends (RACE) was performed as previously described (Frohman 1993) with modifications. Briefly, total RNA was isolated with TRIzol Reagent (Invitrogen) and treated by RNase-Free DNase I (Lucigen) according to the manufacturer’s instructions. The DNase I-treated RNA was extracted by equal volume of RNase-free phenol-chloroform (pH 4.5) and followed by ethanol precipitation by adding 1/10 volume of 3 M sodium acetate (pH 5.3). A sequence-specific oligonucleotide was used to prime the synthesis of first-strand cDNA with a template-switching oligonucleotide (TSO) tail by Superscript II reverse transcriptase (Invitrogen). Three micrograms of RNA were required for setting up reverse transcription reaction (100 U Superscript II reverse transcriptase, 10 U RNase inhibitor, 1 × Superscript II first-strand buffer, 5 mM DTT, 1 M Betaine, 6 mM MgCl_2_, and 1 μM TSO). The reaction was incubated in a thermal cycler with a heated lid (42°C, 90 min; 50°C, 2 min, 42°C, 2 min for 10 cycles; and a final extension at 70°C, 15 min). After the synthesis of first-strand cDNA, PCR amplification was performed with specific primers (98°C, 2 min; 98°C, 10 s, 62.5°C, 30 s, 72°C, 30 s for 32 cycles; and a final extension at 72°C, 2 min) and processed to TA cloning confirmed by Sanger sequencing. Primers for RACE are listed in Supplementary Table S1. All RACE experiments were performed with at least two biological replicates.

#### RNA-seq

RNA-seq experiments were performed as previously described (Guo et al. 2015) with modifications. Briefly, about 10^6^ cells were used for each RNA-seq experiment and total RNA was extracted by TRIzol Reagents (Invitrogen). The polyadenylated mRNAs were enriched from 1 μg of total RNA by the oligo(dT) coupled to magnetic beads (PerkinElmer) and fragmented by heating for 10 min at 95°C. After reverse transcription of the first-strand cDNA and synthesis of the second-strand cDNA, cDNA was purified by 1.8-fold volumes of AMPure XP beads (Beckman). Libraries were constructed with the NEXTflex Rapid RNA-seq Kit (PerkinElmer) according to the manufacturer’s instructions. The purified cDNA was end-repaired and ligated with NEXTflex adapters. The ligated cDNA was amplified by PCR (98°C, 2 min; 98°C, 30 s, 65°C, 30 s, 72°C, 1 min for 14 cycles; and a final extension at 72°C, 4 min). All RNA-seq libraries are sequenced on an Illumina HiSeq X Ten platform.

#### ChIP-seq

Chromatin immunoprecipitation (ChIP) experiments were performed as previously described (Guo et al. 2012) with modifications. Briefly, 4 × 10^6^ HEC-1-B cells were cross-linked by 1% formaldehyde at room temperature for 10 min and quenched by 1 ml of 2 M glycine at a final concentration of 200 mM. Cells were lysed twice with 1 ml of ice-cold ChIP buffer I (10 mM Tris-HCl pH 7.5, 1 mM EDTA, 1% Triton X-100, 0.1% SDS, 0.1% sodium deoxycholate, 0.15 M NaCl, and 1 × protease inhibitors) for 10 min with slow rotations. After centrifuging at 2,500 g for 5 min at 4°C, the lysed cells were resuspended in 700 μl of ice-cold ChIP buffer and sonicated with a high energy setting at a train of 30 s sonication with 30 s interval for 30 cycles using a Bioruptor Sonicator to obtain DNA fragments ranging from 200∼500 bp. After removing the insoluble debris, the cell lysate was pre-cleared with 25 μl of protein A-agarose beads (Millipore) for 2 h at 4°C with slow rotations and immunoprecipitated with an antibody specifically against RNAPII (Millipore), H3K4me3 (Millipore), H3K27ac (Abcam), CTCF (Millipore), or Rad21 (Abcam) rotating slowly overnight at 4°C. The protein-DNA complexes were enriched with 50 μl of protein A-agarose beads (Millipore) by incubating for 3 h at 4°C with slow rotations. After centrifuging at 2,500 g for 1 min at 4°C, the beads were washed once with 1 ml of ChIP buffer I, once with 1 ml of ChIP buffer II (10 mM Tris-HCl pH 7.5, 1 mM EDTA, 1% Triton X-100, 0.1% SDS, 0.1% sodium deoxycholate, 0.4 M NaCl), once with 1 ml of ChIP buffer III (10 mM Tris-HCl pH 7.5, 1 mM EDTA, 1% Triton X-100, 0.1% SDS, 0.1% sodium deoxycholate), and finally with 1 ml of LiCl buffer (0.5 M LiCl, 50 mM Tris-HCl pH 7.5, 1 mM EDTA, 1% NP-40, 0.7% sodium deoxycholate) by incubating for 10 min at 4°C with slow rotations. The cross-linked complexes were then eluted from the thoroughly-washed beads with 400 μl of elution buffer (50 mM Tris-HCl pH 8.0, 10 mM EDTA, 1% SDS) for 1 h at 65°C with constant shaking at 1,000 rpm. The complexes were finally reverse-crosslinked by heating overnight at 65°C. DNA was purified from the reverse-crosslinked complexes with 400 μl of phenol-chloroform, followed by ethanol precipitation by adding 40 μl of 3 M sodium acetate (pH 5.3) and 1,000 μl of ice-cold ethanol. One microliter of glycogen (20 mg/ml) was also added to facilitate DNA precipitation. The purified DNA was end-repaired and then ligated with NEXTflex adapters (PerkinElmer). After PCR amplification (98°C, 2 min; 98°C, 30 s, 65°C, 30 s, 72°C, 1 min for 14 cycles; and a final extension at 72°C, 4 min), all ChIP-seq libraries were sequenced on an Illumina HiSeq X Ten platform.

#### MeDIP-seq

The methylated DNA immunoprecipitation (MeDIP) followed by high-throughput DNA sequencing was performed as previously published (Weber et al. 2005) with modifications. Briefly, 5 × 10^6^ HEC-1-B cells were collected and incubated overnight at 55°C in 500 μl of MeDIP lysis buffer (10 mM Tris-HCl pH 7.4, 2 mM EDTA, 0.2% SDS, 200 mM NaCl) which also include 10 μl of 10 mg/ml proteinase K and 10 μl of 10 mg/ml RNase A. The DNA was purified by phenol-chloroform extraction and ethanol precipitation. The precipitated DNA was dissolved in 100 μl of 10 mM Tris-HCl (pH 7.5). Three micrograms of DNA in a total volume of 100 μl of 10 mM Tris-HCl (pH 7.5) were sonicated with a high energy setting at a train of 30 s sonication with 30 s interval for 18 cycles using a Bioruptor system to obtain DNA fragments below 1,000 bp.

One microgram of sonicated DNA was end-repaired and ligated with NEXTflex adapters (PerkinElmer). The ligated DNA was denatured for 10 min at 95°C and chilled on ice for 10 min because the 5mC (5-methylcytosine) antibody (Active Motif) has a higher affinity for 5mC-containing ssDNA. The ssDNA was diluted in 500 μl of immunoprecipitation buffer (15 mM Tris-HCl pH 7.5, 1 mM EDTA) and incubated with 1 μl of 5mC antibody overnight at 4°C with rotating. The methylated DNA was enriched with 50 μl of protein A-agarose beads (Millipore) by incubating for 3 h at 4°C with slow rotations. After centrifuging at 2,000 g for 1 min at 4°C, the beads were washed thrice with 1 ml of immunoprecipitation buffer for 10 min at 4°C with rotating. The beads were eluted in 200 μl of elution buffer (250 mM NaCl, 1% SDS) for 3 h at 55°C with constant shaking at 1000 rpm. The DNA was purified by phenol-chloroform extraction and ethanol precipitation. The precipitated DNA was amplified by high-fidelity DNA Polymerase (PerkinElmer). All MeDIP-seq libraries were sequenced on an Illumina HiSeq X Ten platform.

#### ChIRP with silver staining

The ChIRP protein assay was performed as previously published (Chu et al. 2015) with modifications. Briefly, 10^9^ cells were used for each silver staining experiment. HEC-1-B cells were cross-linked with 3% formaldehyde (Thermo) for 30 min. The cross-linking reaction was quenched by 1 ml of 1.25 M glycine for 5 min. The cell pellet was harvested by centrifuging at 2,000 g for 5 min at 4°C. The pellet was weighed and dissolved in 1 ml of ChIRP lysis buffer per 100 mg of cell pellet. The cell lysate was sonicated with a high energy setting at a train of 30 s sonication with 30 s interval for 30 cycles using the Bioruptor system to obtain DNA fragments ranging from 1,000 to 2,000 bp. After centrifuging the sonicated cell lysate at 16,000 g for 10 min at 4°C, two-fold volumes of ChIRP hybridization buffer were added into the cell lysate. Pools of biotinylated odd or even *PEARL* probes were added and incubated overnight at 37°C with constant shaking at 90 rpm. The annealed complexes were enriched by streptavidin beads as described in ChIRP-seq.

The beads were resuspended in 100 μl of biotin elution buffer (12.5 mM D-biotin, 7.5 mM HEPES pH 7.5, 75 mM NaCl, 1.5 mM EDTA, 0.15% SDS, 0.075% sarkosyl, and 0.02% sodium deoxycholate) and rotated for 20 min at room temperature, then incubated for 10 min at 65°C. The suspension was transferred into a new tube and the beads were eluted again in 100 μl of biotin elution buffer. Finally, 200 μl of eluents were pooled for protein precipitation. A quarter volume of TCA was added into the eluent and the pulled-down proteins were precipitated by rotating overnight at 4°C. The protein pellet was washed with cold acetone after centrifuging at 16,000 g for 30 min at 4°C. The solubilized pellets were dissolved in 30 μl of 1 × protein loading buffer and denatured for 30 min at 95°C. The denatured proteins were finally separated by SDS-PAGE (sodium dodecyl sulfate-polyacrylamide gel electrophoresis).

The SDS-PAGE gel was stained using a fast silver staining kit (Beyotime) and all steps were performed with constant shaking at 65 rpm at room temperature. Briefly, the gel was fixed in 100 ml of fixing buffer (50 ml of ethanol, 10 ml of acetic acid, and 40 ml of ddH_2_O) for 20 min and washed once with 100 ml of 30% ethanol for 10 min. The gel was washed twice with 200 ml of ddH_2_O for 10 min. The gel was then incubated with 100 ml of silver staining buffer for 10 min and then washed with 100 ml of ddH_2_O for 1.5 min. One hundred milliliters of silver staining coloration buffer were added for 3∼10 min until the desired bands appeared and then 100 ml of silver staining termination buffer were added for 2∼5 min. The stained gel was scanned by PowerLook 2100 XL (UMAX), a high-resolution gel imaging system.

#### Western blot

For the purpose of the total protein extraction, 10^6^ cells were lysed with the RIPA lysis buffer (50 mM Tris-HCl pH 7.4, 150 mM NaCl, 1% Triton X-100, 1% sodium deoxycholate, 0.1% SDS, and 1 × protease inhibitors). Protein lysate was denatured at 95°C for 5 min. Next, proteins were separated by SDS-PAGE and transferred to nitrocellulose membranes. The membranes were blocked by 5% nonfat milk in 1 × TBST solvents and incubated overnight at 4°C with a primary antibody. After washing with 1 × TBST three times, the membranes were incubated at room temperature for 1 h with the secondary antibody and scanned by the Odyssey System (LI-COR Biosciences).

#### Quantitative RT-PCR

About 10^6^ cells were used for each experiment to detect gene expression level by quantitative RT-PCR. Total RNA was extracted by TRIzol Reagents (Invitrogen) and 1 μg of total RNA was used for reverse transcription. The RNA was diluted in 12 μl of nuclease-free water and 4 μl of gDNA Wiper mix (Vazyme) were then added to remove genomic DNA. The reaction was incubated at 42°C for 2 min. Four milliliters of HiScript II Enzyme mix (Vazyme), containing the oligo(dT) and random primers, were added to initiate the reverse transcription reaction. The cDNA was diluted and used for detecting the expression levels of *HS5-1* eRNA *PEARL*, *Pcdhα*, and *GAPDH* by quantitative PCR with the SYBR Green master (Roche). Primers for quantitative RT-PCR are listed in Supplementary Table S1. All quantitative RT-PCR experiments were performed with at least three biological replicates.

### Statistical analysis

Quantitative RT-PCR experiments were performed with at least three biological replicates. RNA-seq, total RNA-seq, ChIP-seq, QHR-4C, ChIRP-seq, MeDIP-seq and DRIP-seq experiments were performed with at least two biological replicates. All statistical tests were performed with GraphPad Prism (v7.0). Data were presented as the mean ± SEM. Statistical significance was determined using unpaired Student’s *t*-test. P ≤ 0.05 was represented as ‘*’. P ≤ 0.01 or 0.001 was represented as ‘**’ or ‘***’, respectively.

### Data Availability

High-throughput sequencing files (RNA-seq, ChIP-seq, QHR-4C, ChIRP-seq, MeDIP-seq, DRIP-seq) have been deposited in NCBI’s Gene Expression Omnibus (GEO) database and are accessible through GEO Series accession number GSE148958. (https://www.ncbi.nlm.nih.gov/geo/query/acc.cgi?acc=GSE148958).

Raw imaging data have been deposited to Mendeley Data: DOI: 10.17632/wfdtgtymry.1 (https://data.mendeley.com/datasets/wfdtgtymry/draft?a=5fb94a1e-8568-41f3-a3c8-06a853d5883f).

*HS5-1* eRNA PEARL sequences have been deposited to Genbank and are accessible through BankIt accession number MZ004931.

---

**Supplemental Fig. S1.**
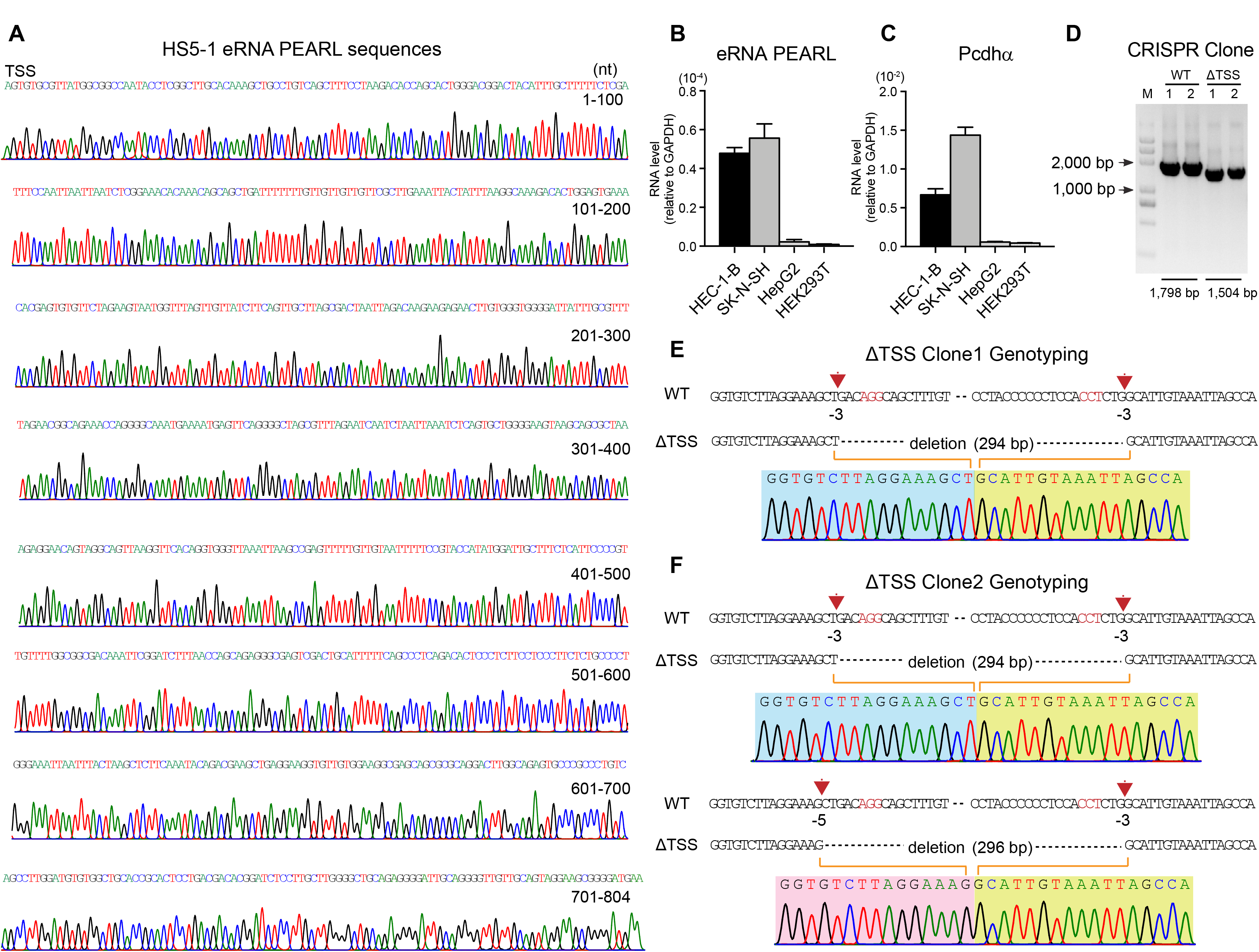
Characterization of the eRNA *PEARL* in the *Pcdhα* Cluster. (A) Sanger sequencing traces of the *HS5-1* eRNA *PEARL* obtained from 5’ RACE experiments. (B-C) The expression levels of the *HS5-1* eRNA *PEARL* and *Pcdhα* in HEC-1-B, SK-N-SH, HepG2, and HEK293T cells. (D) Genotyping of single-cell CRISPR deletion clones. Lane 1: 5 kb DNA ladder. Lane 2-5: PCR products of two 1,798-bp wild-type and two 1,504-bp *HS5-1* eRNA TSS deletion homozygous single-cell CRISPR clones. (E-F) Sanger sequencing traces of the two TSS deletion homozygous single-cell CRISPR clones (ΔTSS clone 1 and ΔTSS clone 2).

**Supplemental Fig. S2.**
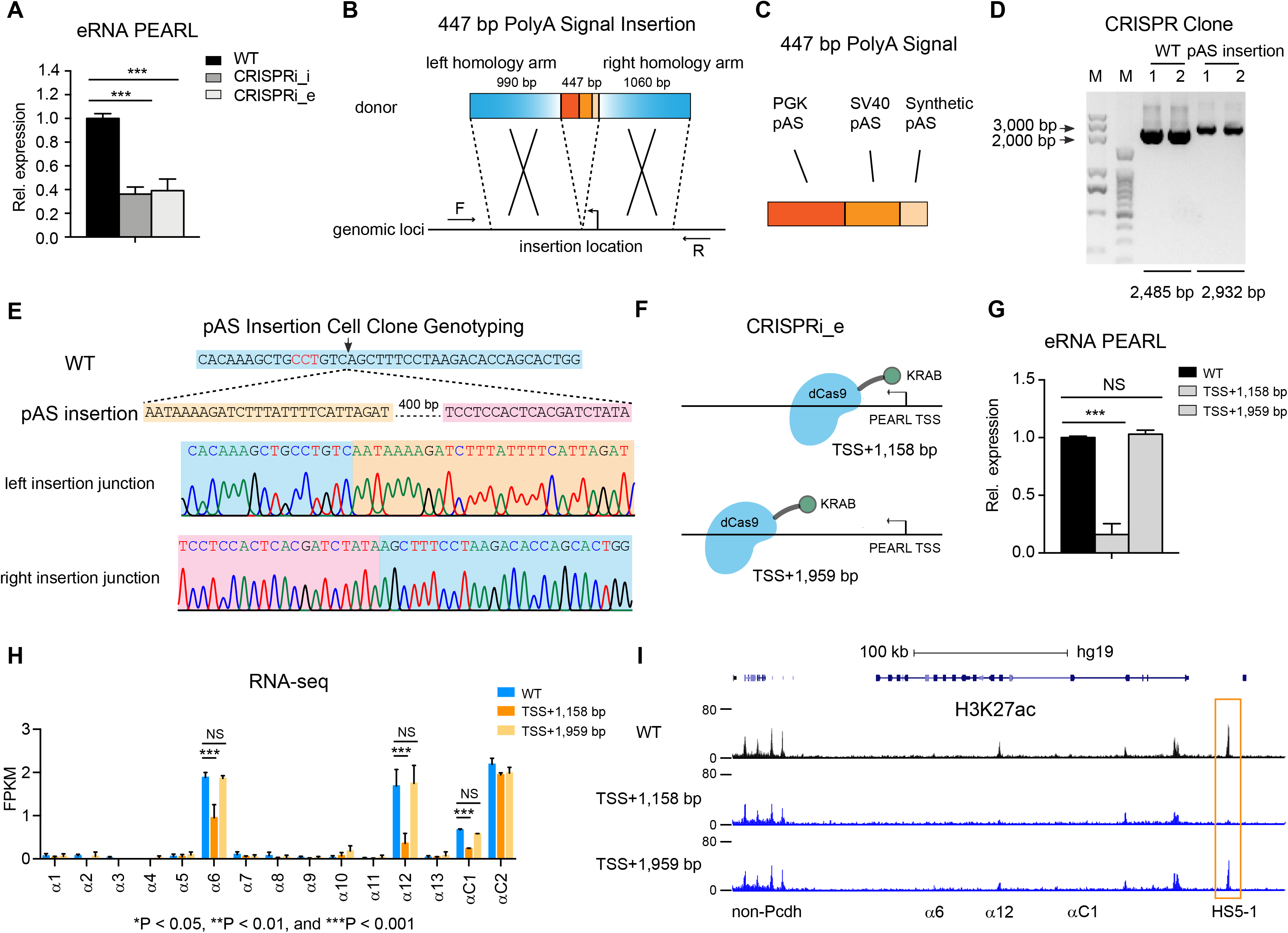
CRISPR Epigenetic and Genetic Perturbation of the *HS5-1* eRNA *PEARL*. (A) Quantitative analyses of eRNA *PEARL* expression after interfering with transcription initiation or elongation (n=3). (B) Schematic of the pAS insertion by homologous recombination using the CRISPR/Cas9 editing system. (C) Schematic of the 447-bp polyadenylation signal sequences including a PGK pAS, a SV40 pAS and a synthetic pAS. (D) Genotyping of the homozygous single-cell CRISPR insertion clones. Lane 1-2: 5 kb and 1.5 kb DNA ladder, respectively. Lane 3-6: Agarose gel electrophoresis for detecting 2,485-bp wild-type bands and 2,932-bp pAS insertion bands of the single-cell CRISPR clones. (E) Sanger sequencing traces of the insertion homozygous single-cell CRISPR clones. (F) Schematic representation of the positions of sgRNAs targeting 1,158-bp and 1,959-bp downstream of the *HS5-1* eRNA *PEARL* TSS. (G) Quantitative analyses of the *HS5-1* eRNA *PEARL* after perturbing transcription elongation by CRISPRi_e with different sgRNAs (n=3). (H) RNA-seq of *Pcdhα* expression levels (n=3). (I) H3K27ac ChIP-seq profiles of the *Pcdhα* and its flanking regions. Data are mean ± SD, *P < 0.05, **P < 0.01, ***P < 0.001; unpaired Student’s *t*-test.

**Supplemental Fig. S3.**
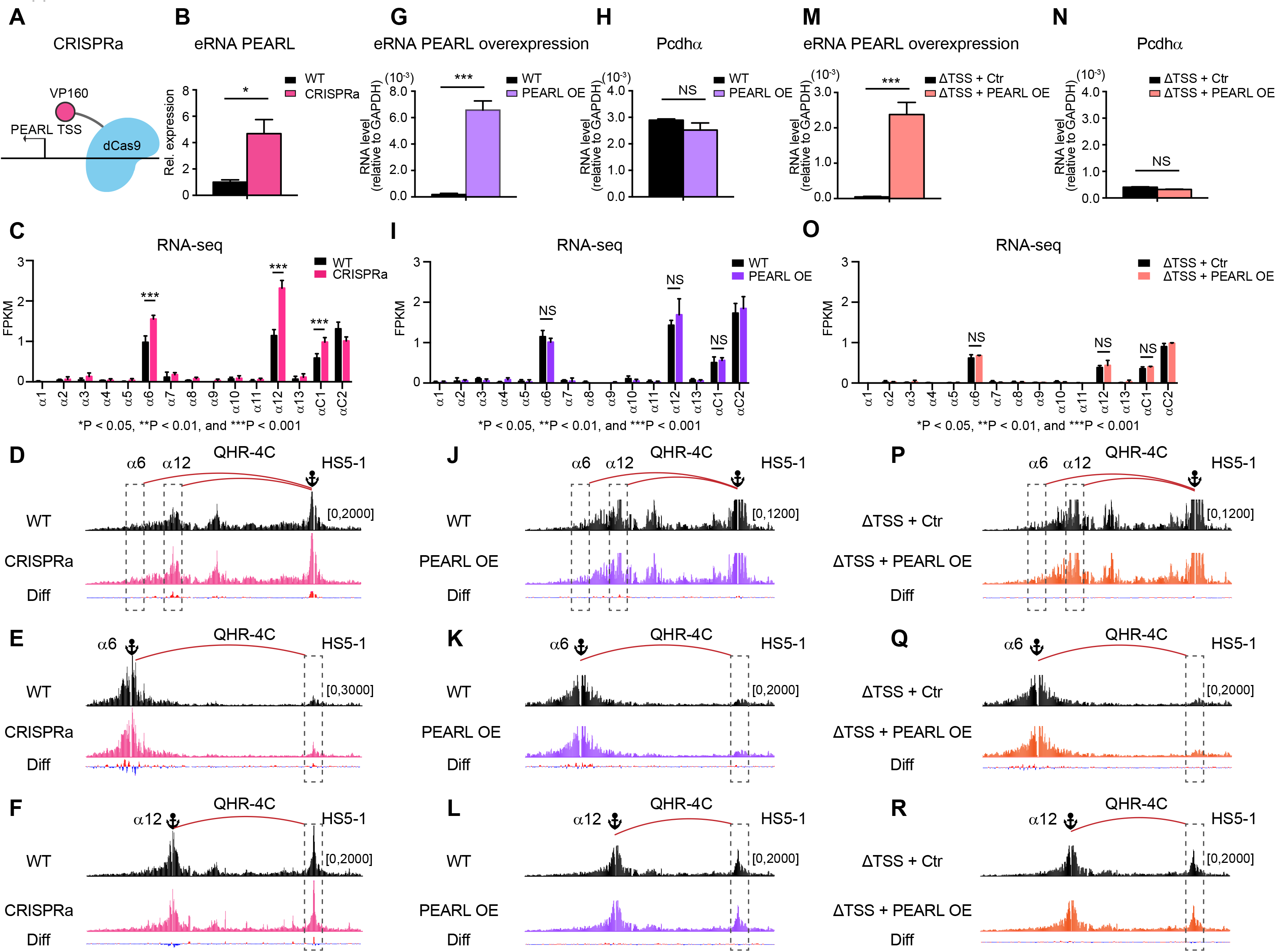
Locally Transcribed *PEARL* Regulates *Pcdhα* Expression. (A) Schematic of CRISPR activation (CRISPRa) with sgRNAs ranging from 361-bp to 557-bp upstream of the *HS5-1* eRNA TSS. (B) The *HS5-1* eRNA expression after activating the *HS5-1* eRNA transcription (n=3). (C) Shown are RNA-seq after activating the *HS5-1* eRNA transcription in the *Pcdhα* cluster (n=3). (D-F) QHR-4C chromatin interaction profiles of WT and CRISPRa with the *HS5-1*, *Pcdhα6,* or *Pcdhα12* as a viewpoint (n=2). (G) Overexpression of the eRNA *PEARL* in HEC-1-B cells (n=3). OE: overexpression. (H) Shown are the *Pcdhα* expression levels after eRNA overexpression (n=3). (I) RNA-seq for the *Pcdhα* cluster after overexpressing the eRNA *PEARL* (n=3). (J-L) QHR-4C chromatin interaction profiles of WT and eRNA overexpression with the *HS5-1*, *Pcdhα6,* or *Pcdhα12* as a viewpoint (n=2). (M-N) Overexpression of the *HS5-1* eRNA *PEARL* cannot rescue *Pcdhα* expression in the ΔTSS clone (n=3). (O) RNA-seq shows no rescue of the ΔTSS clone (n=3). (P-R) QHR-4C chromatin interaction profiles of WT and eRNA overexpression in the ΔTSS clone with the *HS5-1*, *Pcdhα6,* or *Pcdhα12* as a viewpoint (n=2). Data are mean ± SD, *P < 0.05, **P < 0.01, ***P < 0.001; unpaired Student’s *t*-test.

**Supplemental Fig. S4.**
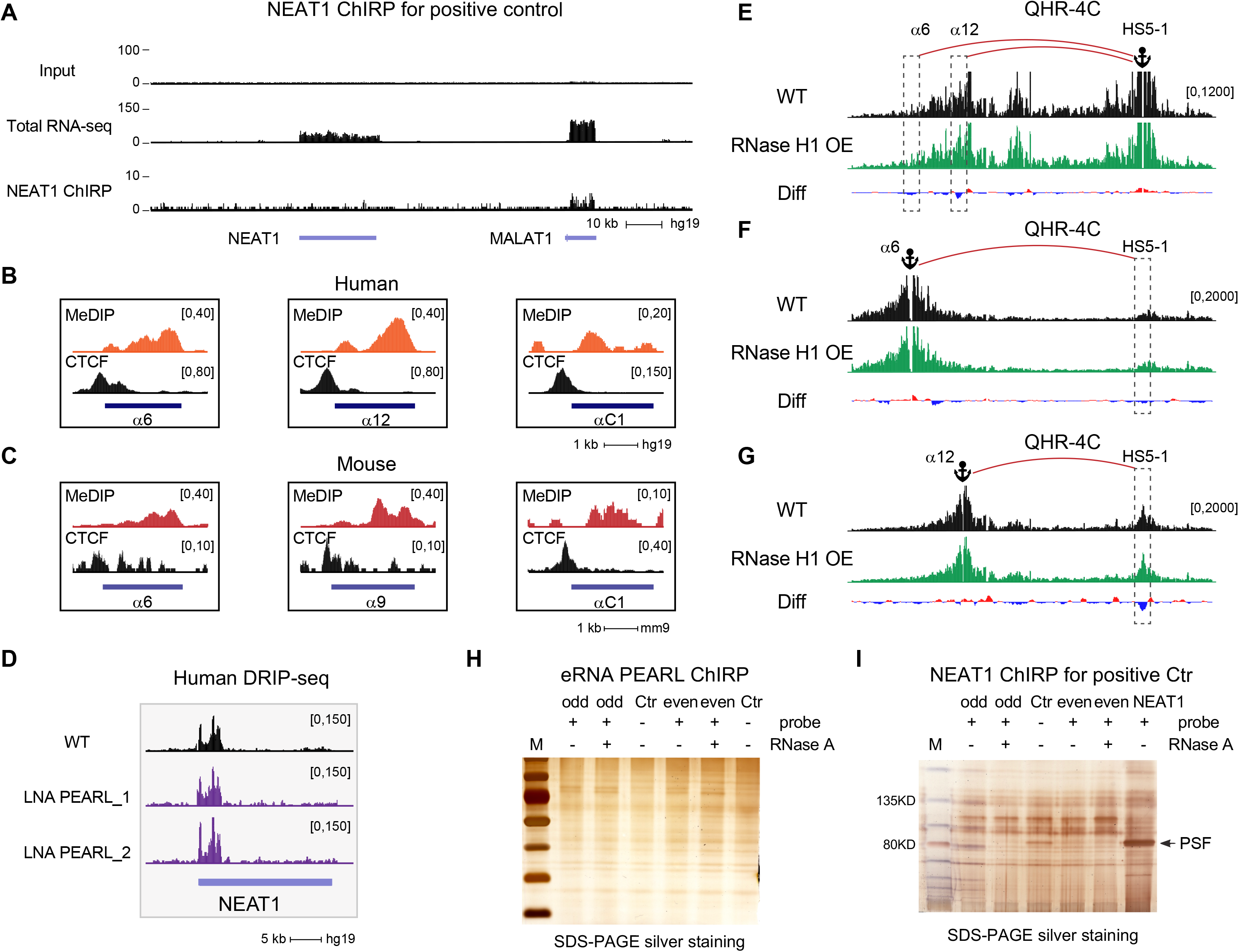
Additional ChIRP, MeDIP, and DRIP Data Supporting *HS5-1* eRNA *PEARL* Regulates Long-distance Chromatin Interactions via R-loop Formation. (A) *NEAT1* ChIRP-seq shows its function *in trans* by heterologous localization at the *MALAT1* locus. (B) The MeDIP-seq experiment at the *Pcdh α6*, *α12,* and *αC1* variable exons in HEC-1-B cells. (C) MeDIP-seq at the *Pcdh α6, α9,* and *αC1* variable exons in mouse cortical tissues. (D) DRIP-seq shows no effect on the *NEAT1* locus after LNA-mediated silencing of the *HS5-1* eRNA *PEARL* transcripts. (E-G) QHR-4C chromatin interaction profiles of WT and RNase H1 overexpression with the *HS5-1*, *Pcdhα6,* or *Pcdhα12* as a viewpoint (n=2). (H) Silver staining of the *HS5-1* eRNA *PEARL* ChIRP pull-down by SDS-PAGE. (I) Identification of *NEAT1* positive ChIRP pull-down proteins by SDS-PAGE with silver staining. Probes (-) and RNase A (+) are served as negative controls. Data are mean ± SD, *P < 0.05, **P < 0.01, ***P < 0.001; unpaired Student’s *t*-test.

## Supplemental Table S1-S2

**Supplemental Table S1.**
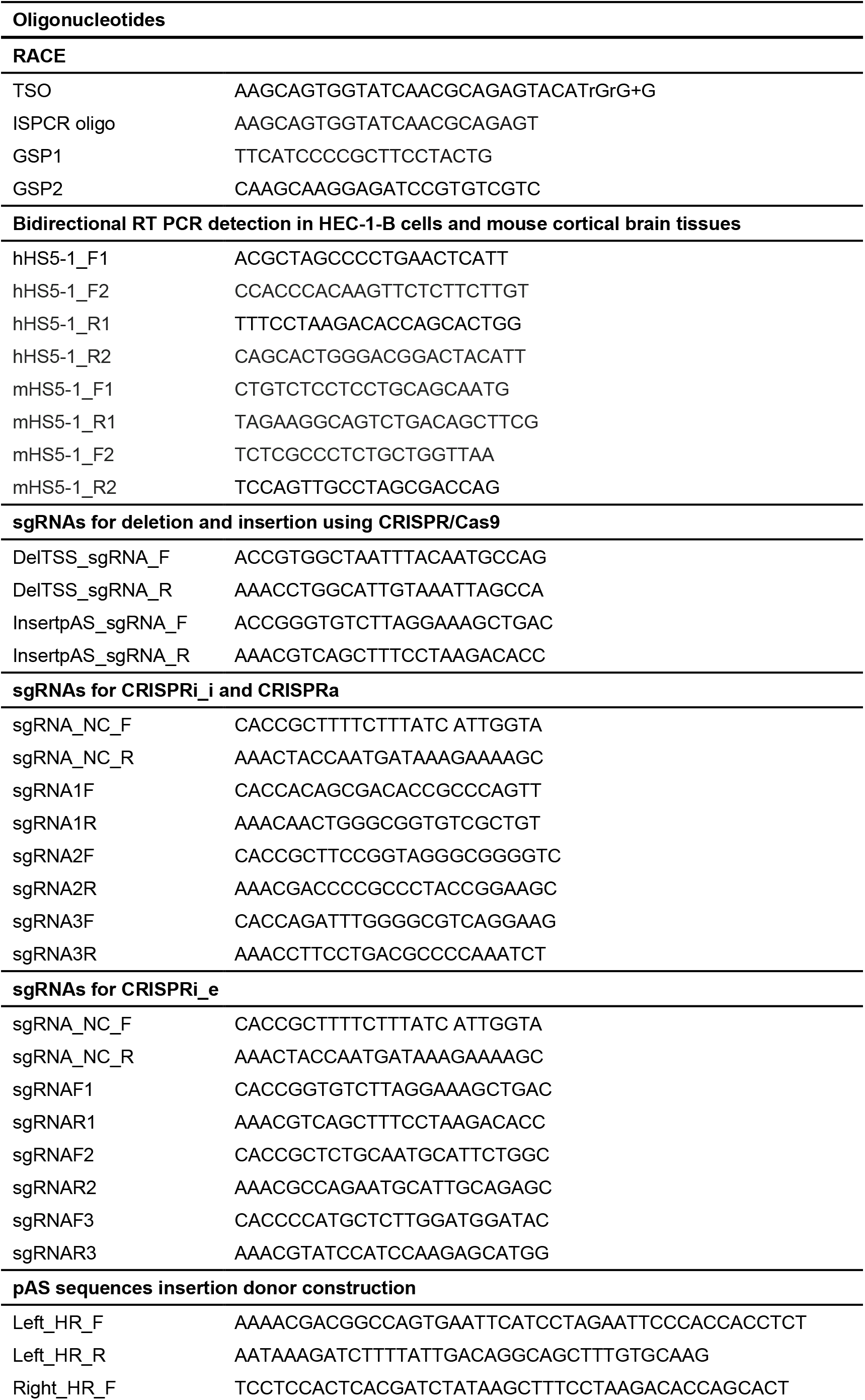
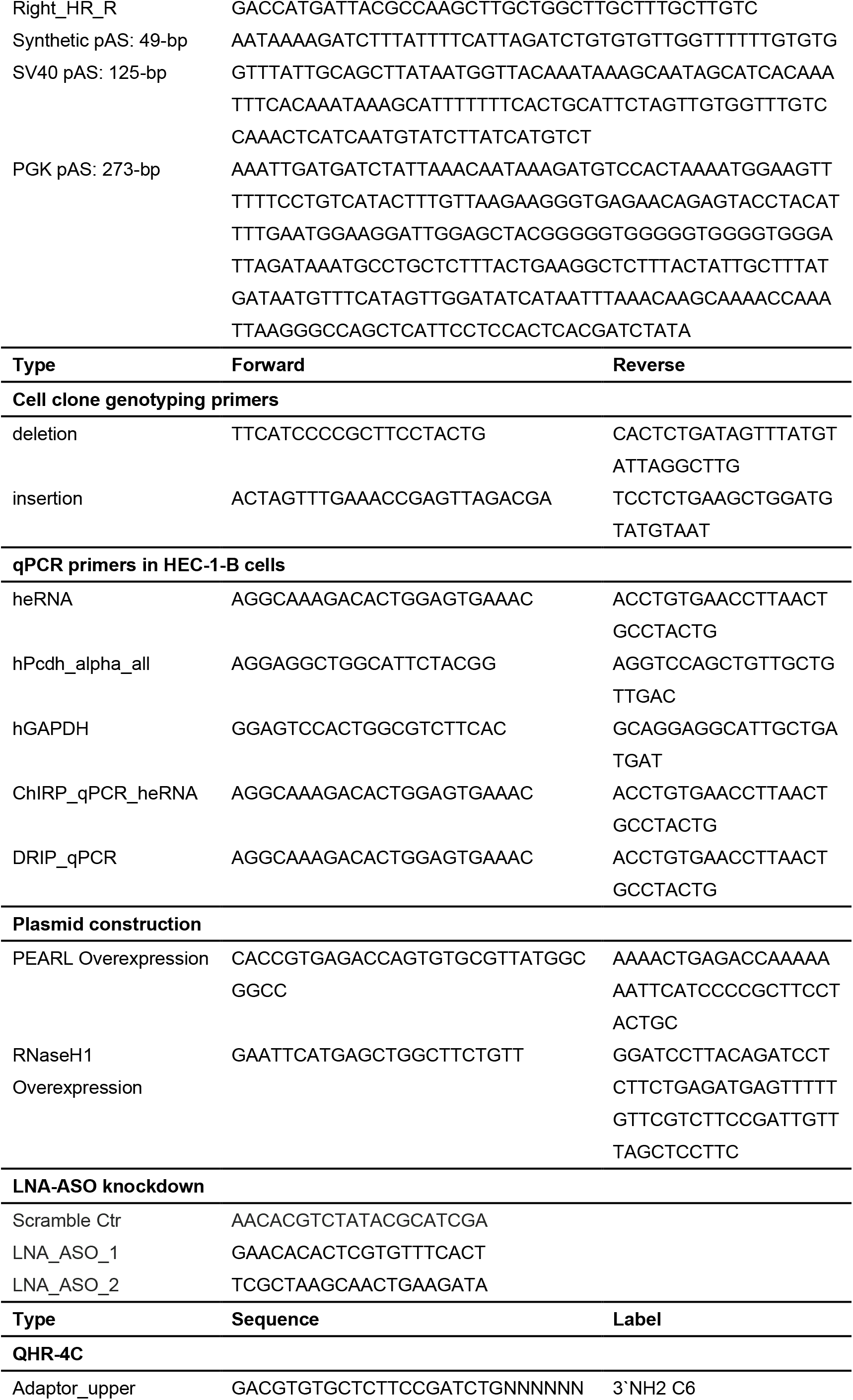
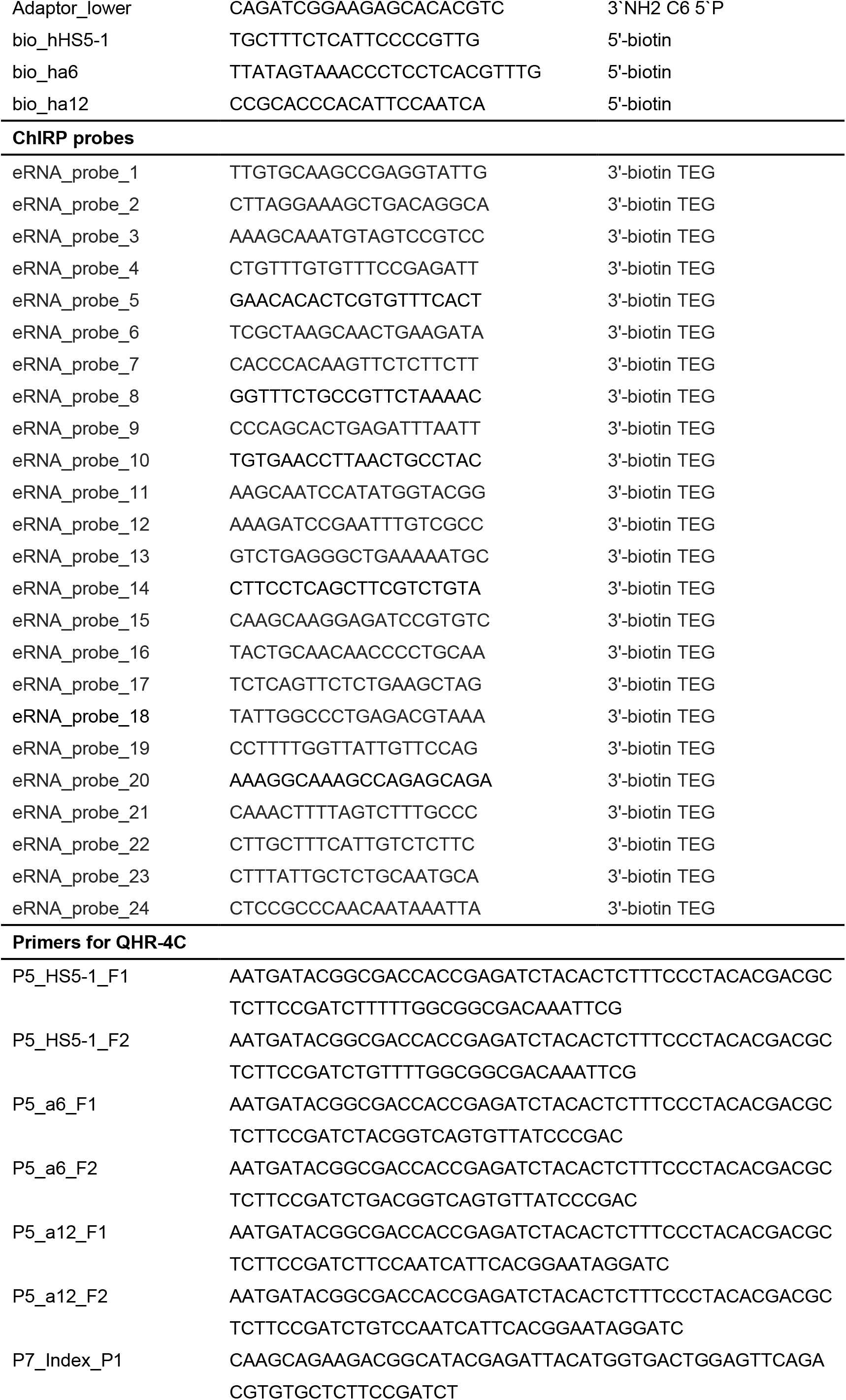
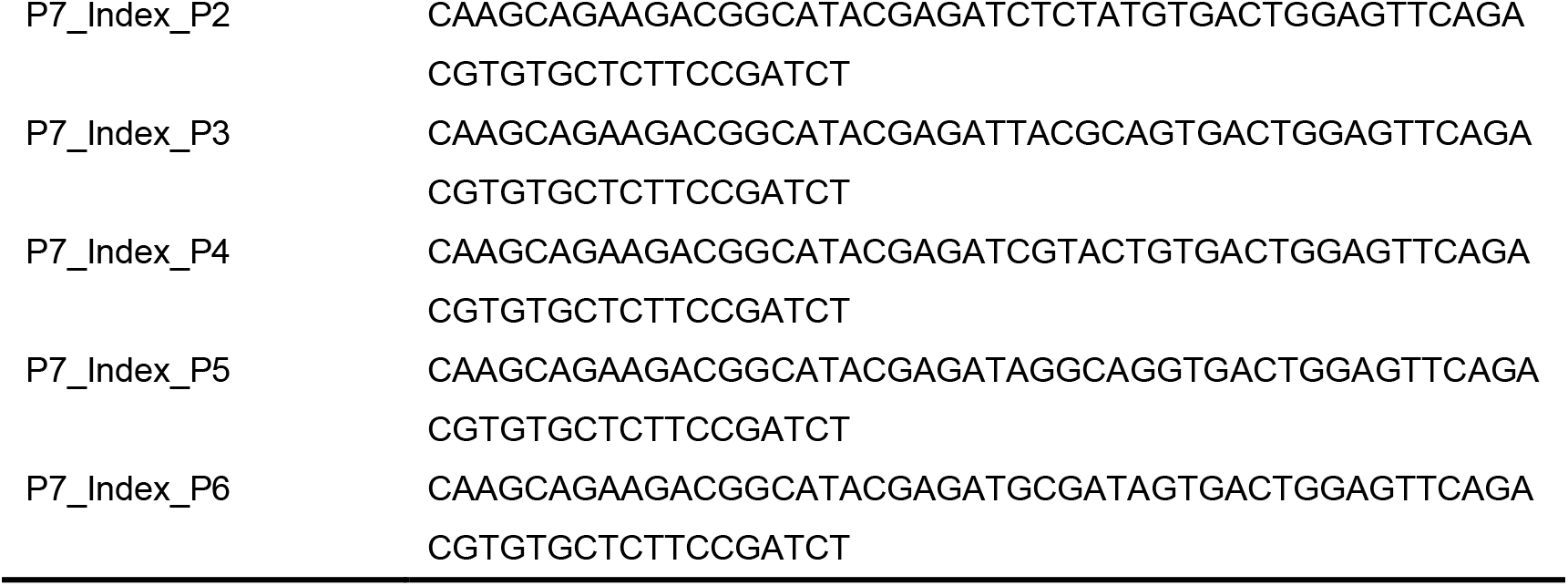
Oligonucleotides used in this study

**Supplemental Table S2.**
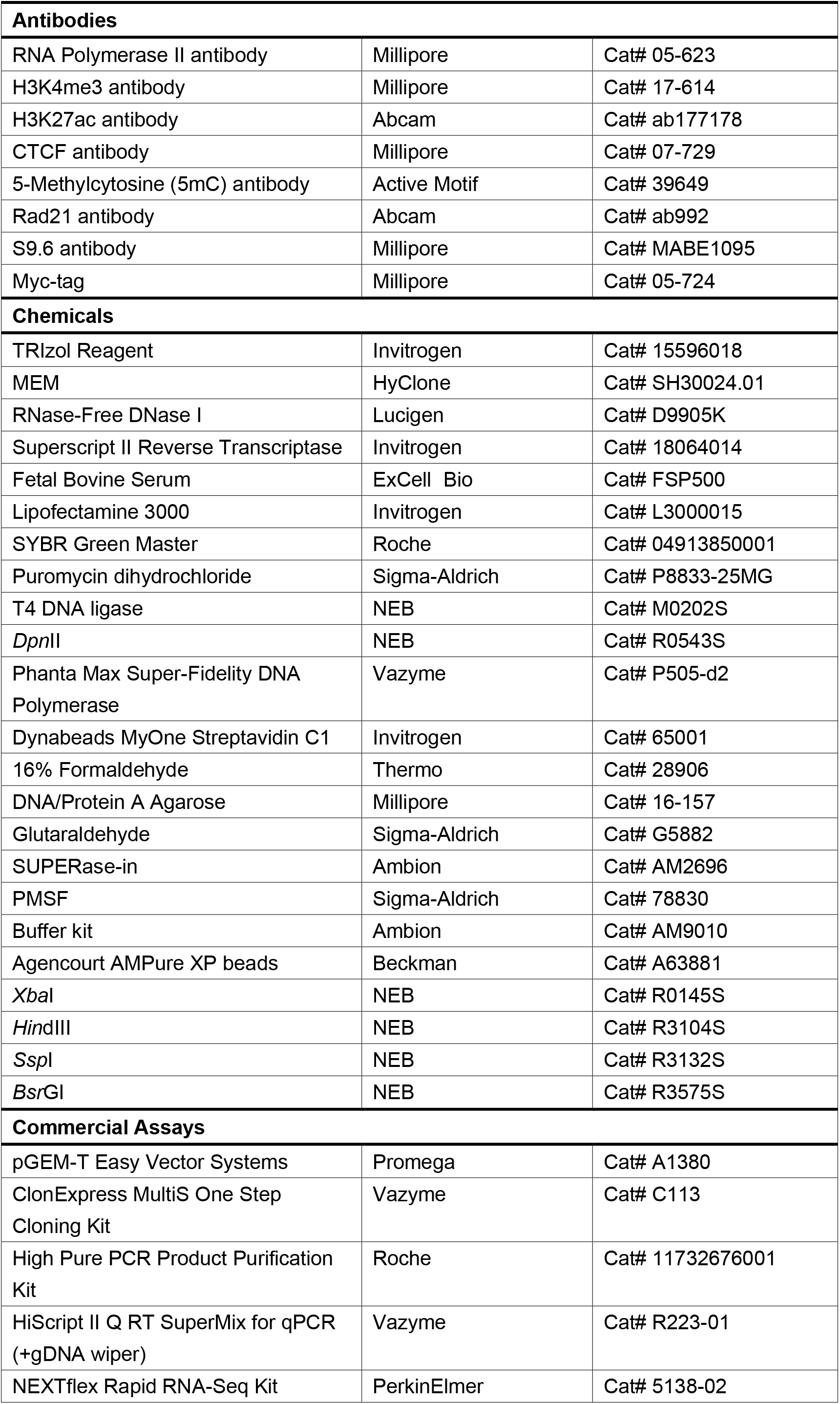
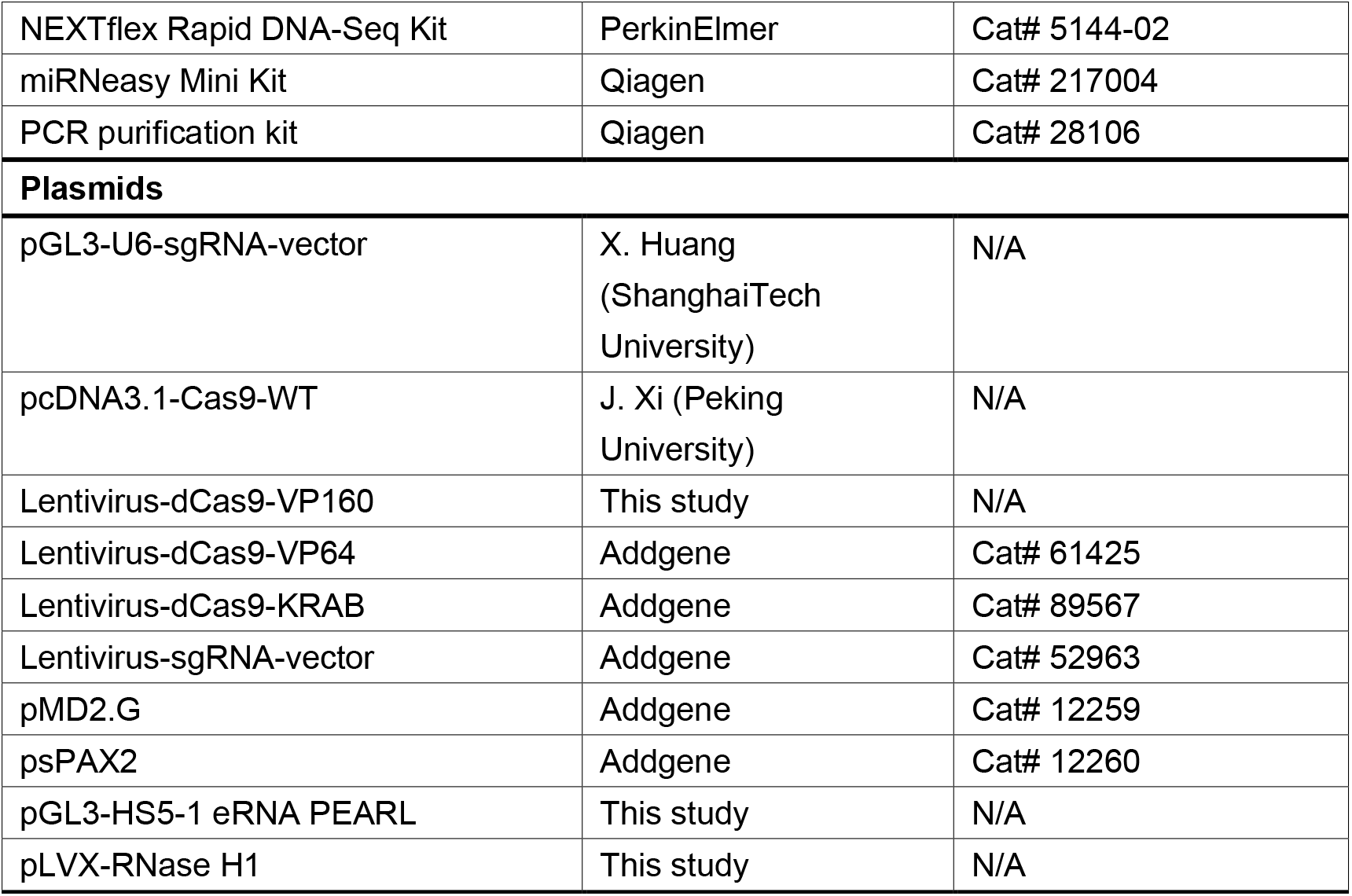
Regents and plasmids used in this study

